# A dynamic gene regulatory code drives synaptic development of hippocampal granule cells

**DOI:** 10.1101/2025.03.27.645686

**Authors:** Blanca Lorente-Echeverría, Danie Daaboul, Jeroen Vandensteen, Gabriele Marcassa, Willem Naert, Joris Vandenbempt, Elke Leysen, Malou Reverendo, Ine Vlaeminck, Lise Vervloessem, Jochen Lamote, Suresh Poovathingal, Kristofer Davie, Keimpe Wierda, Dan Dascenco, Stein Aerts, Joris de Wit

## Abstract

Connecting neurons into functional circuits requires the formation, maturation, and plasticity of synapses. While advances have been made in identifying individual genes regulating synapse development, the molecular programs orchestrating their action during circuit integration of neurons remain poorly understood. Here, we take a multiomic approach to reconstruct gene regulatory networks (GRNs), comprising transcription factors (TFs), regulatory regions, and predicted target genes, in hippocampal granule cells (GCs). We find a dynamic gene regulatory code, with early and late postnatal GRNs regulating cell morphogenesis and synapse organization and plasticity, respectively. Our results predict sequential regulations, with early-active TFs delaying the activation of later GRNs and their putative synaptic targets. Using a loss-of-function approach, we identify *Bcl6* as a regulator of pre- and postsynaptic structural maturation, and *Smad3* as a modulator of inhibitory synaptic transmission, in GCs. Together, these findings highlight the networks of key TFs and target genes orchestrating GC synapse development.

## INTRODUCTION

Neurons are connected into neural circuits via synapses, highly organized and specialized cell-cell junctions. Proper assembly of neural circuits requires neurite outgrowth and guidance, recognition of appropriate cellular partners, and formation of synaptic contacts, followed by maturation and plasticity of synapses (*1–4*). The past decades have seen major progress in the identification of key genes regulating each of these steps in circuit assembly (*5–14*). Understanding how the functions of these individual genes are coordinated to orchestrate the integration of neurons into functional circuits remains elusive.

Gene regulatory networks (GRNs), consisting of transcription factors (TFs), target regulatory regions, and their target genes, coordinate the action of individual genes into molecular programs. Key insight has been derived from the analysis of GRNs in nervous system development of *C. elegans* and *Drosophila* (*15–19*). These studies have postulated the terminal selector hypothesis, in which combinations of constantly expressed TFs define cell identity and key cellular properties such as connectivity. Work in the mouse nervous system has also provided evidence for individual TFs controlling both cellular identity and projection patterns, by regulating expression of axon guidance receptor genes (*20*, *21*). Together, these studies imply a tight link between transcriptional control of cell identity and connectivity pattern. Less is known about the GRNs regulating formation, maturation and plasticity of synaptic contacts. This may be due to complicating factors such as the protracted period required for synapse development in mammalian nervous systems, and the fact that trans-synaptic interactions between pre- and postsynaptic cells play a key role in determining synaptic properties (*22*, *23*).

In this study, we focus on hippocampal granule cells (GCs), a cell type with a well-characterized development in terms of morphology and connectivity. GCs represent the first relay in the hippocampal tri-synaptic loop (Fig. 1A), receiving excitatory input from the entorhinal cortex on their dendrites and providing excitatory input to CA3 pyramidal neurons via their mossy fiber boutons (MFBs), and play an essential role in processing spatial representations and memory acquisition (*24*– *26*). Several studies have examined the role of individual TFs in GC synapse development by loss-of-function approaches (*27–32*). *Sox11* knockout (KO) severely disrupts targeting of GC axons, the mossy fibers (MFs), in the CA3 region and alters MF-CA3 synapse function (*33*). Loss of *Klf9* results in a transient increase in dendritic spine density and length in GCs (*31*). *Mef2c* KO increases dendritic spine density and synaptic transmission at entorhinal cortex-GC synapses (*32*). GCs in *Bcl11b* KO mice show a reduced density of dendritic spines and aberrant MF targeting (*29*). The synaptic organizer C1QL2 (*12*) was recently identified as a target gene of *Bcl11b* (*28*), providing a first causal link between a TF and a synaptic effector in the development of structure and function of GC synapses. However, a comprehensive analysis of the gene regulatory code driving synaptic development of GCs is lacking.

**Fig. 1.**
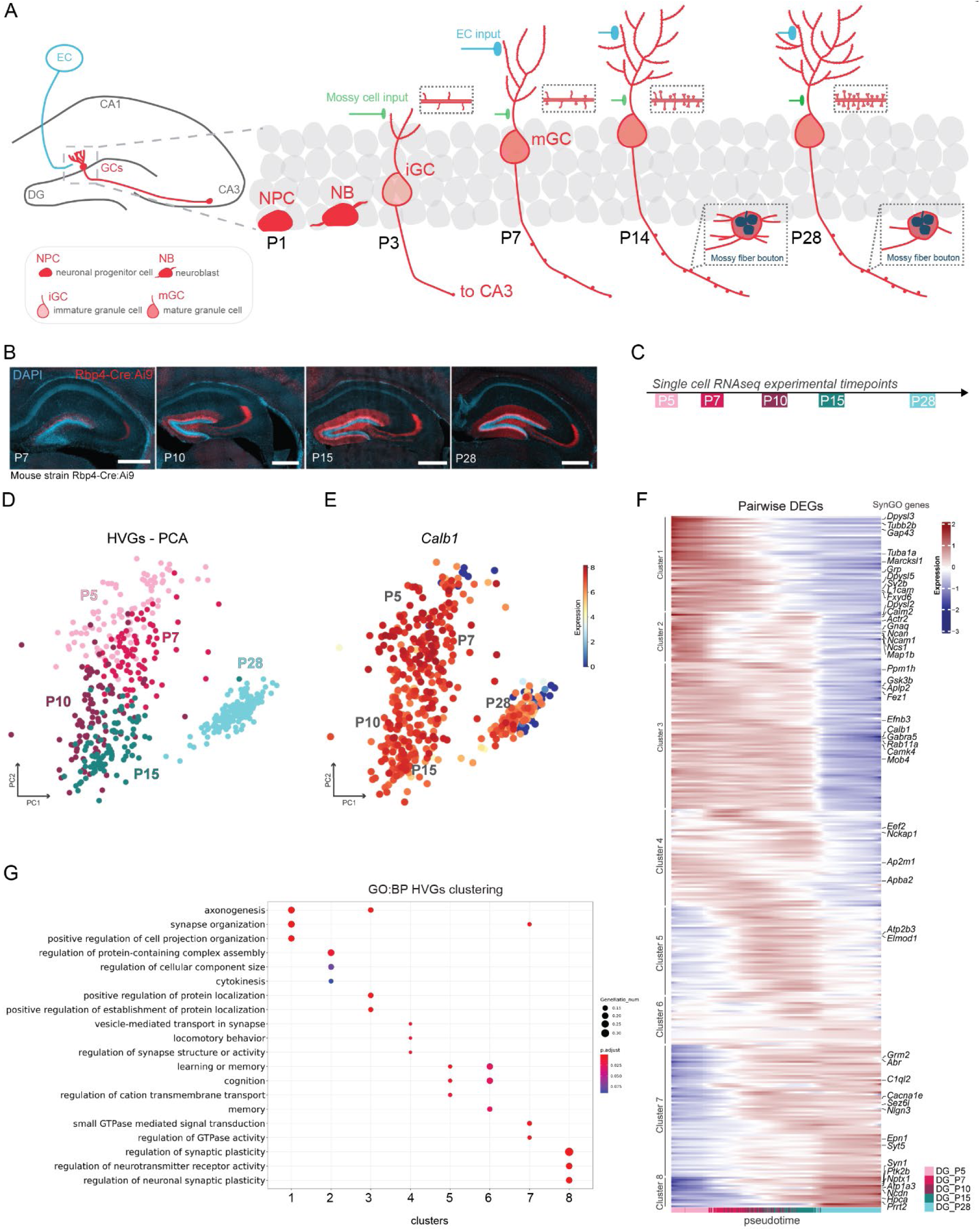
Transcriptomic analysis reveals dynamic changes in synaptic gene expression in GCs. **A)** Scheme of the hippocampal formation and situation of GCs in the hippocampal tri-synaptic circuit: GCs (red) in the dentate gyrus (DG) receive excitatory inputs from the entorhinal cortex (EC, blue) and send axons to CA3. Inset shows postnatal GC development: postnatal day (P)1 NPCs are generated, these will give rise to NBs and further at P3 to iGCs, which extend dendrites to receive input from mossy cells and send axons towards the CA3 region. At P7, iGCs give rise to mGCs, which receive EC input and develop protrusions along their axon. At P14, mGCs present a complex dendritic arbor and increased dendritic spine density, and complex mossy fiber boutons (MFBs) along their axon. At P28, mGCs reach peak dendritic spine density while MFBs are refined, having less filopodia. **B)** Hippocampi of Rbp4-Cre:Ai9 mice at P7, P10, P15 and P28. tdTomato (red) labels GCs in the outer part of the GC layer at each timepoint. Scale bars 500 μm. **C)** Timepoints used for single-cell RNA sequencing experiments. **D)** Principal component analysis (PCA) of highly variable genes (HVGs) separates cells by timepoint. **E)** PCA colored by the expression of GC marker gene Calbindin 1 (*Calb1*). **F)** Heatmap of the eight clusters obtained from the top 50 differentially expressed genes (DEGs) of each pairwise comparison between timepoints, highlighting, for each cluster, genes present in SynGO database within the top 20 genes of each pairwise comparison. Cells are ordered by inferred pseudotime and values represent GAM-fitted expression values for each gene. **G)** Dot plot representing gene ontology (GO) biological process (BP) terms enriched in each cluster.

Here, we first use single-cell RNA sequencing (scRNAseq) to map dynamics in synaptic gene expression in GCs as they integrate into the tri-synaptic hippocampal circuit during postnatal development. We then utilize single-nucleus multiome (snMO) analysis, integrating transcriptome and chromatin accessibility data of the same cells, to reconstruct GRNs. This reveals a dynamic gene regulatory code consisting of early-postnatal GRNs controlling cell morphogenesis and late-postnatal GRNs regulating synapse organization and plasticity. We validate these observations using ribosome tagging, demonstrating that TF and target gene mRNAs are translated in GCs. To test the functional relevance of TFs in GC synapse development, we perform a loss-of-function analysis and identify *Bcl6* as a novel regulator of dendritic spine density and MFB morphology, and *Smad3* as a modulator of inhibitory synaptic transmission. Finally, we construct a GRN model of postnatal GC development based on our results, highlighting how early-postnatal repressors target late-postnatal activators. Together, our findings uncover a dynamic gene regulatory code governing synaptic development of GCs, provide insight in key TFs and target genes, and identify *Bcl6* and *Smad3* as regulators of specific aspects of GC synapse development.

## RESULTS

### Single-cell transcriptomic analysis reveals dynamic changes in synaptic gene expression in GCs

The structural integration of GCs into the hippocampal circuit is well-characterized (Fig. 1A) (*34–40*). In mice, GC neurogenesis peaks around birth, as neural progenitor cells (NPCs) give rise to neuroblasts (NB), which transition into immature GCs (iGCs) around postnatal day (P)3. At this stage, iGCs receive excitatory inputs from mossy cells located in the dentate gyrus (DG) hilus and begin extending axons towards the CA3 region. By P7, iGCs give rise to mature GCs (mGCs), receiving inputs from the entorhinal cortex in the medial molecular layer and displaying some axonal protrusions. By P14, mGCs exhibit an increased dendritic length and spine density, while MFBs in the CA3 region reach peak structural complexity, preceding synaptic remodeling. By P28, mGCs have achieved their final dendritic length and fully developed MFBs. Additionally, GCs start receiving inhibitory input from diverse local GABAergic interneurons during this developmental period (*41*).

To map accompanying changes in gene expression, we performed scRNAseq on GCs from Rbp4-Cre:Ai9 mice at key timepoints during synapse development (Fig. 1B-C). The Rbp4-Cre line specifically expresses Cre recombinase in GCs (*42*, *43*), resulting in tdTomato expression when crossed with Ai9 reporter mice (*44*). We observed that labeled GCs in Rbp4-Cre:Ai9 mice at each developmental time point were located in the outer part of the GC layer, representing more mature GCs (Fig. 1B). tdTomato-positive GCs were fluorescently sorted and processed for single-cell sequencing using SMART-Seq2 to detect a high number of genes per cell (*45*) (Fig. S1A-D and Methods). Principal component analysis (PCA) demonstrated a clear age-based separation of cells (Fig. 1D). Sorted cells expressed the postmitotic GC marker gene *Calb1* (Fig. 1E), as well as other GC and neuronal markers (*Prox1*, *Rfx3*, *Slc17a7*, *Gad1*; Fig. S1E) (*46*, *47*).

Pairwise differential gene expression analysis (MAST test, p-value <0.05, |logFC| > 0.5) revealed significant gene expression changes in GCs across postnatal development, with prominent shifts at P7 and P15 (Fig. S1F). Subsequent clustering of the top 50 differentially expressed genes (DEGs) per pairwise test (see Methods) showed eight clusters according to their gene expression profiles across pseudotime (Fig. 1F). We next examined synaptic gene expression patterns over pseudotime, by analyzing DEGs annotated in the expert-curated SynGO database (*48*) within these clusters and linking them to the function of each cluster using gene ontology (GO) analysis (Biological Process, p-val 0.05, q-val 0.05) (Fig. 1G). In the early clusters (clusters 1-2-3), we found genes related to axonogenesis and regulation of protein-containing complex assembly, like *Gap43* and *Ncam1*, respectively; as well as *Dpysl2* and *Dpysl3*, cytosolic phosphoproteins involved in growth cone collapse and other critical functions in developing neurons (*49*) (Fig. 1F, G). Cluster 3 contained, among others, *Efnb3*, an axon guidance cue (*50*) (Fig. 1F, G). The middle clusters (clusters 4-5-6) contained genes linked to vesicle-mediated transport (e.g. *Apba2* and *Ap2m1*), and others linked to learning and memory. Finally, the late clusters (clusters 7-8) contained genes peaking in expression around P15-28, which are involved in synaptic organization, such as the postsynaptic cell adhesion molecule *Nlgn3*, and synaptic plasticity, such as the secreted synaptic organizer *Nptx1* (Fig. 1F, G).

Together, this analysis reveals the dynamics in synaptic gene expression during circuit integration of GCs, with an early-postnatal (P5-P7) phase related to cell morphogenesis and a late-postnatal phase (P15-P28) regulating synapse organization and plasticity.

### Single-nuclei multiome analysis reveals GRN dynamics during postnatal development of GCs

To map the GRNs underlying the dynamic changes observed in synaptic gene expression during postnatal GC development (Fig. 1F, G), we generated a 10X snMO dataset of P5, P10, P15 and P28 wildtype mouse hippocampi (Fig. 2A, B). Combined transcriptome and chromatin accessibility from the same nuclei enables the reconstruction of enhancer-driven regulons (eRegulons) comprising TFs, their target regulatory regions and target genes (*51*). After sequencing and quality controls, cell types were annotated, and GC lineage nuclei were extracted using marker genes (Fig. S2A-H and Methods) (*46*, *47*). NPCs, found in the subgranular zone of the DG, are the most immature cells, identified based on *Cdk6* and *Top2a* expression (Fig. S2A). They are chronologically followed by type 1 NBs, expressing *Elavl2* and *Calb2*, and type 2 NBs, which express *Rgs6*. Type 2 NBs give rise to iGCs, marked by *Pcdh15* and *Prox1* expression, and finally mGCs, the main excitatory neurons of the DG, expressing *Meg3*, *Calb1*, and *Ntng1* (Fig. S2A). Marker genes (Fig. S2A) and similarity analyses using the cosine metric (Fig. S2B) confirmed that the mGCs in this dataset align best with the GCs from our scRNAseq dataset (Fig. 1), labeled by Rbp4-Cre:Ai9. Each age showed varying proportions in cell types, with a higher proportion of NPCs at P5 and more mGCs at P28 (Fig. S2I). Uniform manifold approximation and projection (UMAPs) based on transcriptome and epigenome profiles depict the transition from NPCs to mGCs (Fig. S2J-L).

**Fig. 2.**
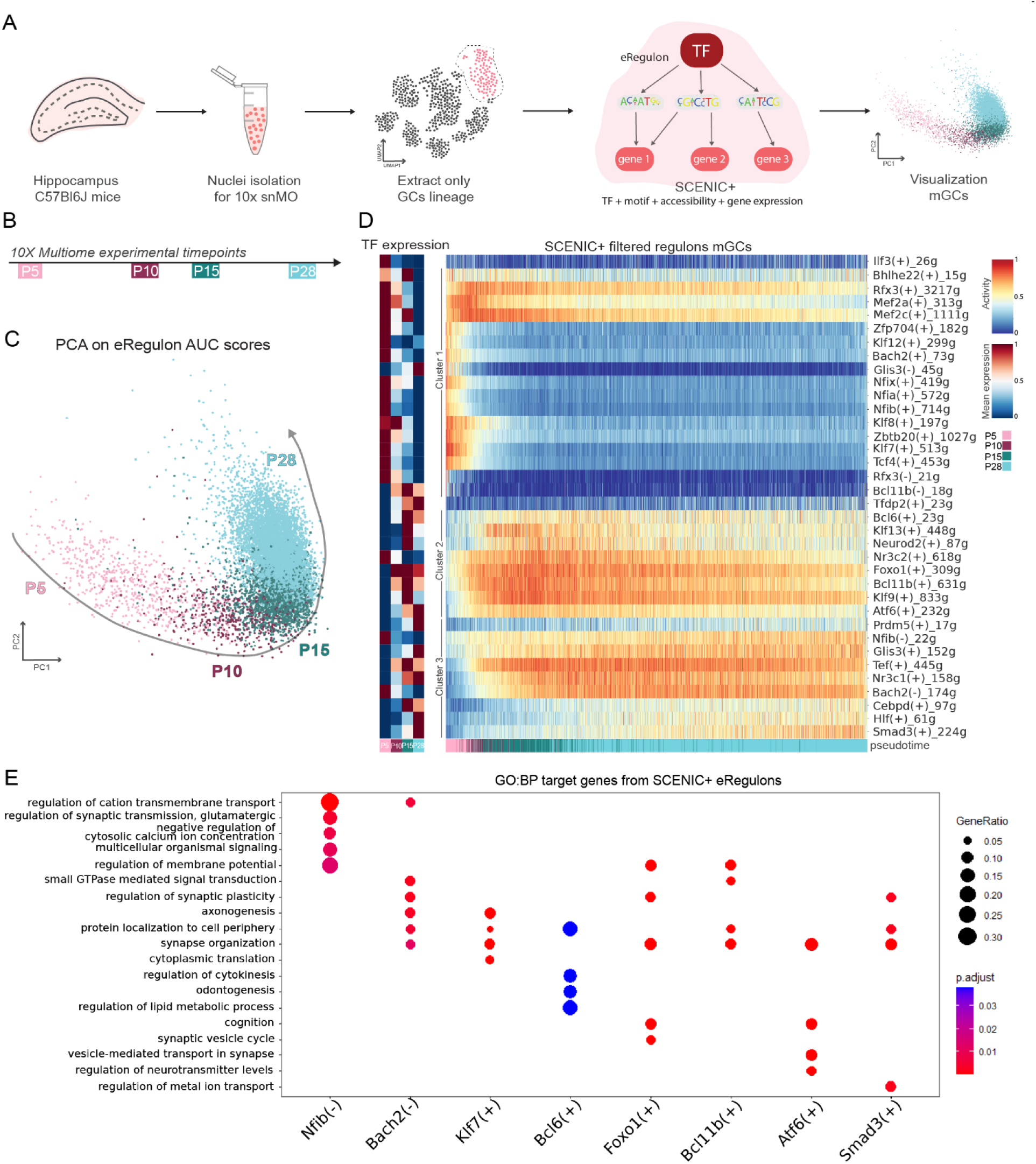
Single-nuclei multiome (snMO) analysis reveals gene regulatory network (GRN) dynamics during postnatal GC development. **A)** Methodology for snMO experiments. Hippocampi from C57Bl6 mice were homogenized, nuclei were isolated and processed for both RNA sequencing and ATAC sequencing. From this dataset the GC lineage was extracted using markers and SCENIC+ was applied to infer eRegulons (transcription factor (TF) – accessible target regions – target genes). Only mature GCs (mGCs) are shown in the following panels. **B)** Timepoints used for snMO experiments. **C)** PCA based on eRegulon AUC (Area Under recovery Curve) scores of target genes separates mGC nuclei by timepoint. **D)** Left panel: heatmap of leading TF expression across time in snMO RNAseq. Right panel: heatmap of SCENIC+ filtered eRegulons’ (see methods for filtering criteria) AUC scores in mGCs across timepoints. Cells are ordered by inferred pseudotime values. **E)** Dot plot representing gene ontology (GO) biological process (BP) terms enriched in the target genes of representative eRegulons.

Focusing on mGCs, we analyzed scATACseq (assay for transposase-accessible chromatin) data to track chromatin accessibility changes over time (Fig. S3A). We focused on pairwise differentially accessible regions (DARs) across consecutive developmental stages, meaning that for each age, we compared its accessible regions with those of the subsequent age (Wilcoxon test, p.adj < 0.01 and log(FC) > 1). By using a simple approach and assigning individual peaks to the closest genes, we observed that DARs are associated with genes with distinct roles at different stages (Fig. S3B). Between P5-P10, DARs associated with genes related to cell fate commitment decrease in accessibility, while regions assigned with genes linked to axonogenesis or synapse organization increase in accessibility. From P15-P28, the accessibility of regions near axonogenesis-related genes decreases, while accessibility for those near genes related to vesicle transport and signal transduction increases. These findings align with the timing of GC maturation and synapse development (Fig. 1A, refs (*34–40*)).

We next applied SCENIC+ (*51*) to the entire GC lineage (see Methods and Fig. S3C-F), and focused on eRegulons active in mGCs across postnatal development (Fig. 2A). PCA analysis on SCENIC+ eRegulons enrichment scores in mGCs (AUC (area under the recovery curve) scores, calculated by AUCell (*52*)) reveals a clear temporal separation from P5 to P28 (Fig. 2C). eRegulon AUC scores represent the enrichment of their target genes within each cell’s expressed genes. A total of 36 eRegulons with distinct activity patterns were identified in mGCs (Fig. 2D, Table S1), with leading TF expression following a similar pattern as eRegulon activity over time for activators, and the opposite pattern for repressors (Fig. 2D, Fig. S3F). eRegulons are defined by the name of their leading TF, and a sign corresponding to their activator or repressor role (i.e. TF(+) or TF(-)), followed by the number of target genes they contain.

eRegulons grouped by activity patterns fell into three main clusters (hierarchical clustering, 0.85 cut-off value, Fig. S3G). The first cluster (cluster 1, early, Fig. S3G, Fig. 2D) was highly active at P5 and included Zbtb20(+) and Klf7(+). The second cluster (cluster 2, middle, Fig. S3G, Fig. 2D) peaked at P10-P15 and decreased thereafter, including Bcl11b(+), Foxo1(+), Bcl6(+) and Atf6(+). The third cluster (cluster 3, late, Fig. S3G, Fig. 2D) peaked at P15-P28 and included Smad3(+) and the repressors Bach2(-) and Nfib(-), among others.

We next explored whether eRegulon activity patterns during synaptic development of mGCs correspond to target gene function, using GO analysis (Biological Process). Klf7(+) targets are associated with axonogenesis (Fig. 2E), consistent with *Klf7*’s role in promoting axon growth (Table S1) (*53*). Bcl11b(+) regulates target genes involved in membrane potential and synapse organization (Fig. 2E), in agreement with the role of the BCL11B TF in the DG (Table S1) (*28*). Smad3(+) targets genes are important for synaptic plasticity and ion transport, aligning with Smad3’s role in long-term potentiation (LTP) in GCs (*54*) (Fig. 2E, Table S1). Target genes of the *Nfib* repressor are involved in synaptic transmission and membrane potential (Fig. 2E), consistent with the fact that their expression is repressed at P5. Similarly, Bach2(-) targets genes are related to plasticity and synapse organization. Of the 18 eRegulons without a known role in the DG (Table S1), Bcl6(+) targets are related to protein localization to cell periphery, and Atf6(+) controls genes involved in vesicle transport at the synapse (Fig. 2E). Taken together, these findings reveal a dynamic gene regulatory code, consisting of known and novel GRNs, orchestrating postnatal development of mGCs.

### Predicted TFs and their targets are translated in developing GCs

We next determined whether the identified TFs and target genes are translated in GCs, given the complex relationship between cellular transcriptome and proteome (*55*). To this end, we utilized ribosome tagging to analyze transcripts of TFs and target genes recruited for translation and correlated these with inferences from our snMO analysis. We crossed the RiboTag mouse, which expresses the HA-tagged ribosomal subunit *Rpl22* in the presence of Cre recombinase (*56*), with the Rbp4-Cre line to capture ribosome-bound mRNA in GCs across postnatal stages P5-P28 (Fig. 3A-B). Following hippocampal dissociation, ribosome-bound mRNA was extracted and subjected to bulk RNA sequencing after quality control (Fig. 3C, Fig. S4A-C). PCA showed a clear temporal separation between the samples from P5 to P28, with the P21 and P28 stages being nearly indistinguishable (Fig. 3D). Differential gene expression analysis (pairwise analysis, adj. p-value < 0.05, |log2FC| > 0.5) again revealed prominent shifts at P7 and P15 (Fig. S4C; compare to Fig. S1F), with 5 clusters containing SynGO annotated genes (Fig. S4E) (*48*).

**Fig 3.**
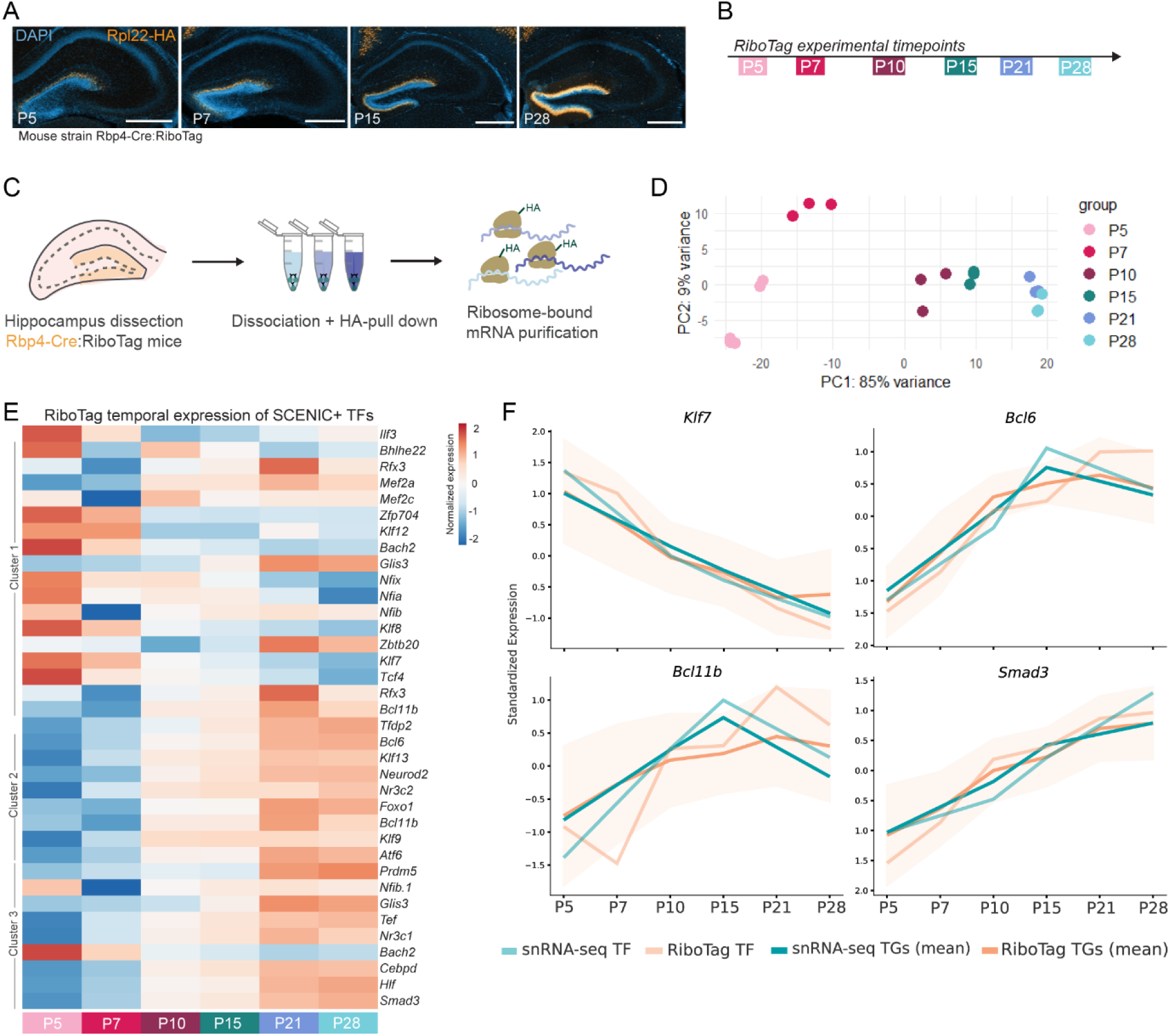
Predicted transcription factors (TFs) and their target genes (TGs) are translated in developing GCs. **A)** Hippocampi of Rbp4-Cre:RiboTag mice at P5, P7, P15 and P28. HA tag (orange) labels the ribosomal subunit RPL22 expressed in GCs in the outer part of the GC layer at all timepoints. Scale bars 500 μm. **B)** Timepoints used for RiboTag experiments. **C)** Methodology for RiboTag experiments. Hippocampi from Rbp4-Cre:RiboTag mice were homogenized and the homogenate was incubated with anti-HA magnetic beads for RPL22-HA pull down. Ribosome-bound mRNA was separated from ribosomes, RNA was isolated and libraries prepared for bulk RNA sequencing. **D)** PCA shows separation of biological replicates by timepoint. **E)** Heatmap of SCENIC+ eRegulon leading TF (compare with Fig. 2D) expression in RiboTag samples across timepoints. Label on the left side of the heatmap indicates hierarchical clustering of the eRegulons (see Fig. S3G). **F)** Comparative plots of standardized expression across time for *Klf7*, *Bcl11b*, *Bcl6* and *Smad3* show TF expression in snMO (gene expression) RNAseq (light blue line), TF expression in RiboTag (light orange line), TGs mean expression in snMO (gene expression) (turquoise line), TGs mean expression in RiboTag (orange line), and TGs mean expression in RiboTag ± 1 standard deviation (cream shade around TGs mean expression).

Using this dataset, we analyzed the expression patterns of ribosome-bound transcripts of TFs leading the SCENIC+ predicted eRegulons in mGCs (Fig. 2D). Ribosome-bound mRNA levels for these TFs largely correlated with their expression patterns in the snMO dataset (Fig. 3E, Fig. S4F). For example, TFs such as *Nfib* or *Klf7* peaked at P5-7 and decreased thereafter, while others, including *Bcl6*, *Atf6*, and *Smad3*, exhibited sharp increases after P15 (Fig. 3E, Fig. S4F). Similarly, TFs with previously characterized roles in development of GC connectivity, *Bcl11b* and *Klf9*, showed matching translatome profiles and expression patterns in the snMO dataset (Fig. 3E, Fig. S4F).

We next assessed eRegulon target genes in the RiboTag dataset. By analyzing mean expression of target genes over time, we observed a similar expression profile between ribosome-bound transcript level and expression pattern in the snMO dataset (Fig. S4G). For example, ribosome-bound mRNA levels for Klf7(+), Bcl11b(+), Smad3(+) and Bcl6(+) target genes (Fig. 3F, orange lines) correlated with their expression pattern in the snMO dataset (Fig. 3F, turquoise lines; Fig. S4G). Thus, the ribosome-bound TF and target gene transcript dynamics show a strong correlation with the predicted eRegulon activity patterns, increasing confidence in the identified GRNs that regulate postnatal development of GCs.

### Loss of *Bcl6* in GCs alters dendritic spine density and MFB morphology

To test the functional relevance of identified GRNs in synaptic development of GCs, we focused on eRegulons from middle and late clusters (clusters 2 and 3; see Fig. 2D, Fig. S3G), as their activity coincides with the timing of formation, refinement and maturation of GC synapses. We selected two GRNs, Bcl6(+) and Smad3(+), based on the roles of their predicted target genes and the unknown role of their leading TFs in GC synapse development. Focusing on these, we performed a loss-of-function analysis of the corresponding TFs in combination with sparse, bright labeling of GCs for morphological analysis.

We first investigated Bcl6(+), an eRegulon from cluster 2 (middle) whose activity peaks at P15 (Fig. 4A). *Bcl6* itself has not been previously linked to synapse development and does not have a known role in the DG. In the cortex, *Bcl6* promotes neurogenesis via repression of Notch and Wnt pathways (*57*, *58*). In the DG, *Bcl6* is not detected in progenitors and iGCs but is expressed in mGCs (*29*). In Rbp4-Cre:Ai9 mice, we detected *Bcl6* mRNA in GCs at P15, but not at P5, using single-molecule Fluorescent *In Situ* Hybridization (smFISH) (Fig. S5A-B), consistent with Bcl6(+) eRegulon activity (Fig. 4A). Bcl6(+) includes 23 target genes encoding proteins with predominantly synaptic localization (Fig. 4B) that have been linked to hippocampal synapse development (*59*, *60*). One of these predicted *Bcl6* target genes is *Nptx1*, encoding a secreted synaptic organizer (*61*) that is highly enriched at MF-CA3 synapses (*9*) and regulates clustering of AMPA receptors (*62*). *Nptx1* is controlled by a chromatin region that peaks in accessibility at P28 (Fig. 4C-D, Fig. S5C).

**Fig 4.**
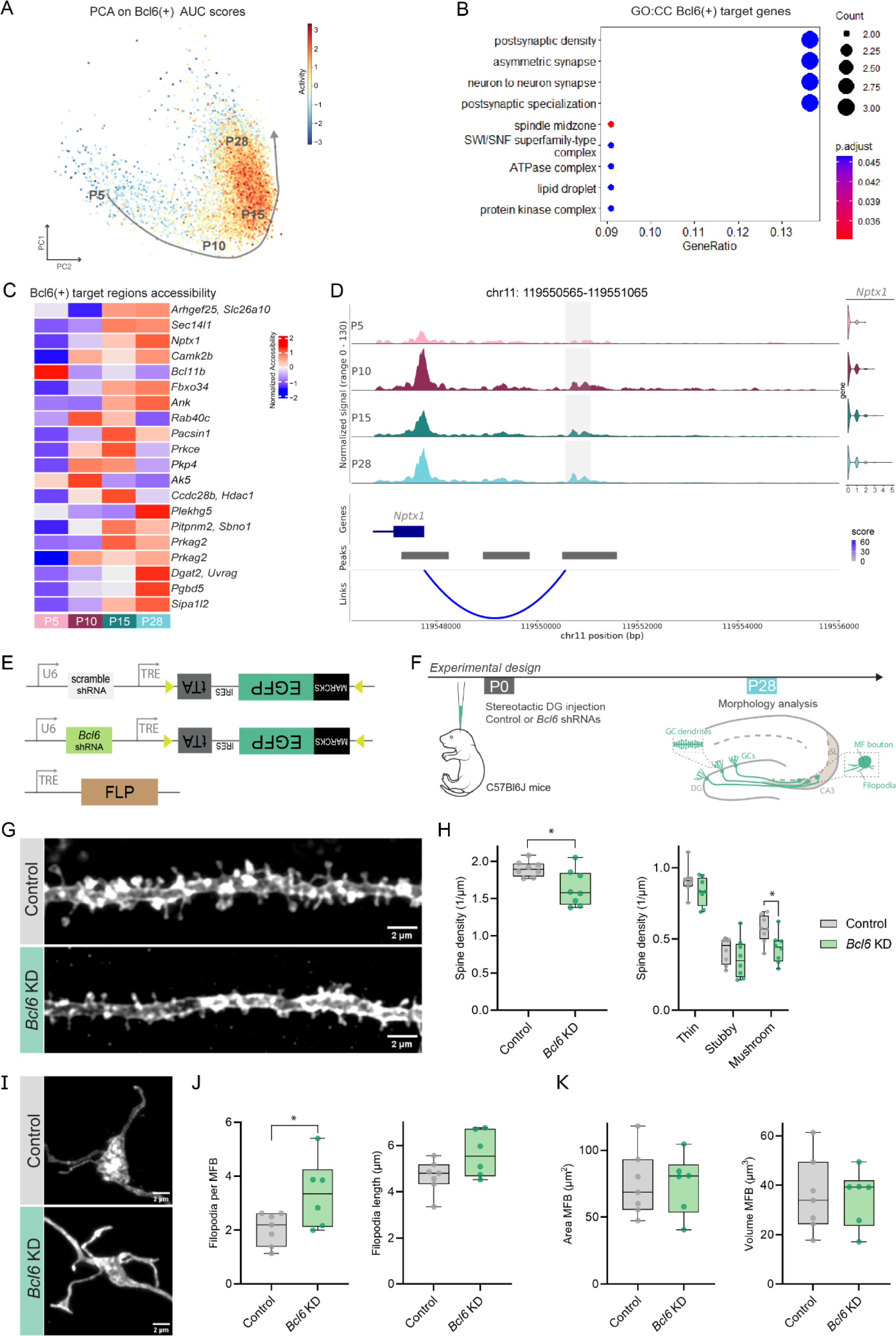
Loss of *Bcl6* in granule cells (GCs) alters dendritic spine density and mossy fiber bouton (MFB) morphology. **A)** PCA based on eRegulon target gene cell enrichment scores (AUC) colored by Bcl6(+) AUC scores of target genes. **B)** Dot plot represents gene ontology (GO) cellular component (CC) enriched terms for the target genes of Bcl6(+). **C)** Heatmap showing mean normalized accessibility per timepoint of Bcl6(+) 24 predicted target regions. **D)** Coverage plot for *Nptx1* associated region across timepoints in mature GCs (mGCs). Target region is located on chromosome (chr) 11 between base pairs (bp) 119550565 and 119551065 (grey shade). Arcs represent SCENIC+ predicted region-gene links. **E)** Plasmids pAAV-U6-TRE-fDIO-mGFP-IRES-tTA containing scrambled shRNA or *Bcl6* shRNA, and pAAV-TRE-FLP. TRE: tetracyclin responsive element, IRES: internal ribosomal entry site, tTA: tetracycline transactivator, MARCKS: myristoylated alanine-rich protein kinase C substrate. FRT (FLP recognition target) sites are indicated with yellow arrowheads. **F)** Experimental design for shRNA experiments. C57Bl6J pups were injected in the DG with control or *Bcl6* shRNA AAVs and AAV-TRE-FLP for sparse labeling at P0 and perfused at P28 for morphological analysis of GC dendrites (spine density and morphology) and MFB morphology (area, volume, filopodia number and length). **G)** Representative images of dendritic segments of GCs from the medial molecular layer of the DG for control (grey) and *Bcl6* KD animals (green). **H)** Spine density is reduced in *Bcl6* KD animals (n=4 20 µm dendrite segments analyzed per animal, n=8 mice for control; n=8 mice for *Bcl6* KD; nested t-test, p-value = 0.0135). Mushroom spine density is significantly reduced (nested t-test, p-value = 0.0220) in *Bcl6* KD animals compared to controls. **I)** Representative images of MFBs from control (grey) or *Bcl6* KD animals (green). **J)** MFB filopodia number is increased in *Bcl6* KD animals compared to controls (6-8 MFBs analyzed per animal; n=7 mice for control; n=6 mice for Bcl6 KD; nested t-test, p-value = 0.0209), while filopodia length is unchanged. **K)** MFB area and volume are unchanged upon *Bcl6* KD (6-8 MFBs analyzed per animal; n=7 mice for control; n=6 mice for *Bcl6* KD; nested t-test). Box-and-whisker plots in H, J and K show median, interquartile range, minimum, and maximum; with each dot representing one animal.

To explore *Bcl6*’s role in synaptic development of GCs, we used short hairpin RNA (shRNA)-mediated knockdown (KD) of *Bcl6,* along with a flippase (FLPo) for Supernova-mediated sparse labeling (Fig. 4E, S5D) (*63*). Attempts to validate *Bcl6* guide RNAs (gRNAs) for CRISPR/Cas9-mediated knockout (KO) were unsuccessful due to a lack of specific BCL6 antibodies. We therefore resorted to previously validated *Bcl6* shRNAs (*57*, *58*), whose efficiency we confirmed in primary neurons (Fig. S5E-F). We injected AAVs expressing either scrambled or *Bcl6* shRNA combined with FLPo (Fig. 4E) into the DG of C57Bl/6J mice at P0. At P28, GC dendritic and axonal segments were imaged and reconstructed in 3D for morphological analysis (Fig. 4F, Fig. S5G). *Bcl6* KD resulted in a 20% decrease in dendritic spine density compared to GCs expressing the scrambled shRNA (Fig. 4G-H). This decrease was caused by a reduction in mature, mushroom spine density (Fig. 4H).

We next analyzed MFB area, volume and filopodia number and length. MFBs contain one or multiple filopodia that mediate feedforward inhibition to CA3 neurons via synapses onto inhibitory neurons (*64*). *Bcl6* KD resulted in a 64% increase in the number of filopodia per MFB, without affecting filopodia length (Fig. 4I-J). MFB area and volume remained unchanged in *Bcl6* KD animals (Fig. 4I, K). Together, these results show that loss of *Bcl6* in GCs reduces dendritic mushroom spine density and increases MFB filopodia density.

### Loss of *Smad3* in GCs alters inhibitory synaptic transmission

We next focused on the Smad3(+) eRegulon, a cluster 3 (late) eRegulon whose activity peaks at P28 (Fig. 5A). Consistent with this, we detected *Smad3* mRNA using smFISH in GCs of Rbp4-Cre:Ai9 mice at P15 and P28, but not at P5 (Fig. S6A). *Smad3* is predicted to regulate 224 genes involved in synaptic plasticity and organization (*10*, *65*, *66*) (Fig. 2E) through 649 chromatin-accessible regions, which gradually increase in accessibility from P5 to P28 (Fig. 5B). One of these predicted *Smad3* target genes is *Tanc1*, a scaffold protein, highly expressed in the hippocampus, that interacts with the excitatory postsynaptic scaffold PSD-95. Its deficiency causes reduced spine density in the CA3 region of the hippocampus (*65*). *Tanc1* is controlled by a chromatin region that peaks in accessibility at P28 (Fig. 5B-C, Fig. S6C).

**Figure 5.**
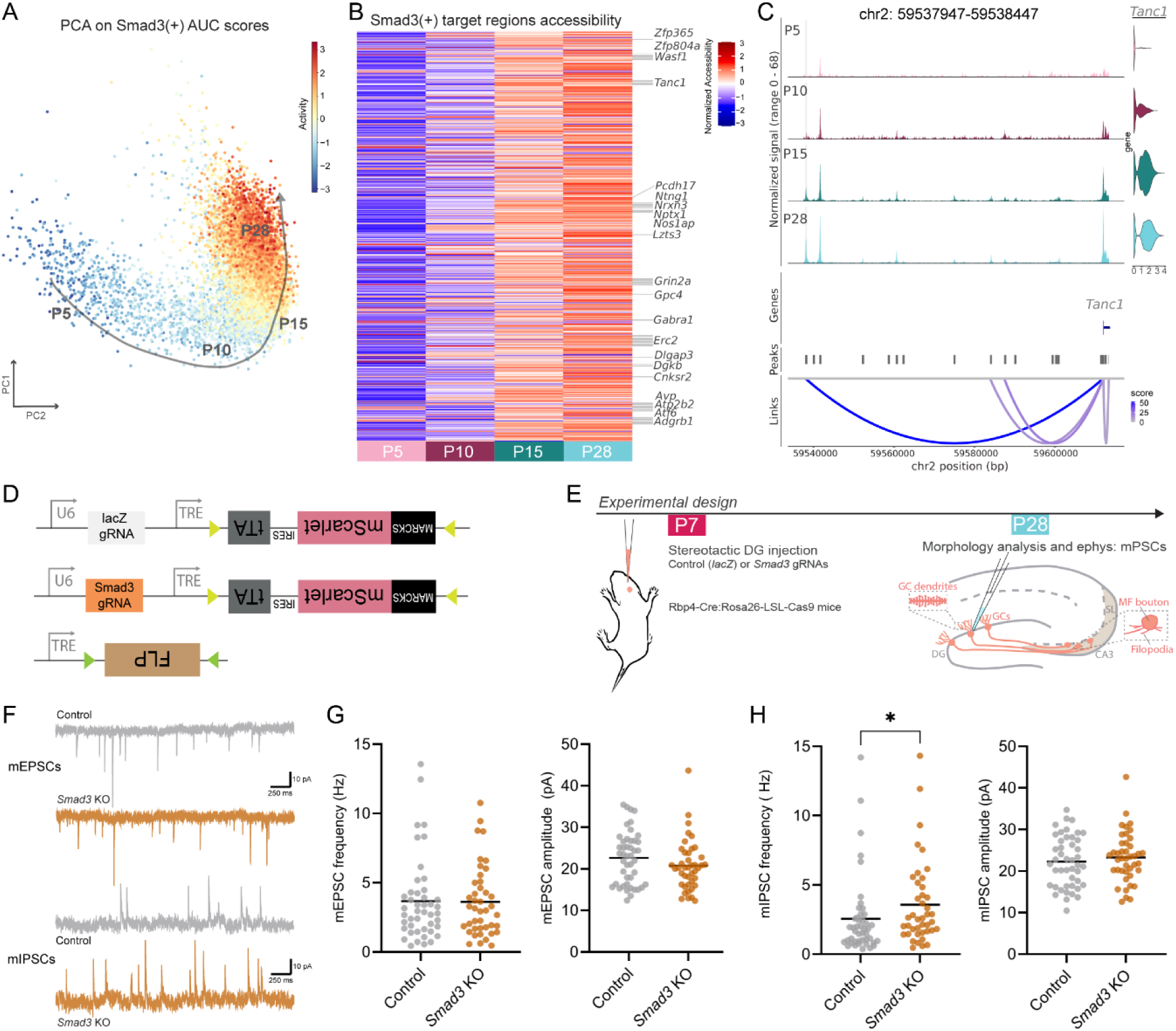
Loss of *Smad3* in granule cells (GCs) alters inhibitory synaptic transmission. **A)** PCA based on eRegulon target gene cell enrichment scores (AUC) colored by Smad3(+) AUC scores of target genes. **B)** Heatmap showing mean normalized accessibility per timepoint of Smad3(+) 649 predicted target regions and labelling region-linked genes within the synaptic organization gene ontology term. **C)** Coverage plot for *Tanc1* associated region across timepoints in mature GCs (mGCs). Target region is located on chromosome (chr) 2 between base pairs (bp) 59537947 and 59538447 (grey shade). Arcs represent SCENIC+ predicted region-gene links. **D)** Plasmids pAAV-U6-TRE-fDIO-mScarlet-IRES-tTA containing control gRNA (lacZ) or *Smad3* gRNAs, and pAAV-TRE-DIO-FLP. FRT sites (FLP recognition target) are indicated with yellow arrowheads; loxP sites (Cre recognition target) are indicated with green arrowheads. **E)** Experimental design for *Smad3* in vivo experiments. Rbp4-Cre:Rosa26-LSL-Cas9 P7 pups were injected in the DG with control or *Smad3* gRNA AAVs and AAV-DIO-TRE-FLP for sparse labeling. Morphological analysis of GC dendrites (spine density and morphology) and MFB morphology (area, volume, filopodia number and length) was performed at P28; electrophysiology at P27-31. **F)** Representative traces of miniature excitatory (mEPSCs) and inhibitory (mIPSCs) postsynaptic currents recorded from GCs expressing control or *Smad3* gRNAs. **G)** Analyzed mEPSC frequency and amplitude (number of neurons n=45 from n=4 mice for control and n=42 from n=4 mice for *Smad3* KO conditions, Mann-Whitney test). **H)** Analyzed mIPSC frequency and amplitude (number of neurons n=45 from n=4 mice for control and n=42 from n=4 mice for *Smad3* KO conditions, Mann-Whitney test, p-value = 0.01).

To assess whether loss of *Smad3* affects GC dendritic spine density or MFB morphology, we used CRISPR/Cas9-mediated KO. *In vitro* validation confirmed that *Smad3* gRNAs effectively reduced SMAD3 protein levels in primary hippocampal cultures (Fig. S6E-F). We crossed Rbp4-Cre mice with Cre-dependent Cas9 mice (Rosa26-LSL-Cas9 (*67*)) for GC-specific *Smad3* KO, combined with an AAV-DIO-FLPo to achieve Supernova-mediated sparse labeling (Fig. 5D-E, Fig. S6D) (*63*). We injected AAVs expressing *Smad3*-targeting gRNAs or a non-targeting (lacZ) gRNA as a control (Fig. 5D-E, S6G) into the DG of Rbp4-Cre:Rosa26-LSL-Cas9 mice at P7 and analyzed GC morphology at P28 (Fig. 5E). Density and morphology of dendritic spines in *Smad3* KO GCs did not differ from control GCs (Fig. S6H, I). Additionally, analysis of area, volume and filopodia number and length of MFBs did not show significant changes following *Smad3* loss (Fig. S6J-L), indicating that *Smad3* KO in GCs does not affect dendritic spine density or MFB morphology.

Loss of *Smad3* has previously been shown to abolish long-term potentiation (LTP) in GCs via enhanced GABA-A receptor neurotransmission (*54*). We therefore extended our analysis and recorded spontaneous synaptic transmission in GCs, using whole-cell voltage-clamp recordings in acute hippocampal slices from Rbp4-Cre:Rosa26-LSL-Cas9 mice (P27-31) injected with AAVs expressing *Smad3* or lacZ gRNAs (Fig. 5D-E). The frequency and amplitude of miniature excitatory postsynaptic currents (mEPSCs) in *Smad3* KO cells did not differ from controls (Fig. 5F-G). In contrast, the frequency of miniature inhibitory postsynaptic currents (mIPSCs) was increased in *Smad3* KO cells compared to controls (Mann-Whitney test, p-value = 0.01), while mIPSC amplitude remained unchanged (Fig. 5F, H).Taken together, these results show that loss of *Smad3* in GCs increases the frequency of spontaneous inhibitory synaptic transmission, without affecting spontaneous excitatory synaptic transmission, dendritic spine density or MFB morphology.

### Early-active GRNs repress late TFs and predicted targets involved in synaptic organization

We next explored how the identified GRNs interact with each other. Most of the leading TFs target other TFs (Fig. S7), belonging to the same or different clusters (see clustering analysis Fig. S3G). For example, cluster 1 TFs (Fig. 2D), such as the Nfi TFs (*Nfib*, *Nfia*, *Nfix*), are highly interconnected, but are also predicted to target later TFs, as seen for Rfx3(+), targeting *Glis3* or *Nr3c1*, among others (Fig. S7).

Focusing on Bcl6(+) and Smad3(+) predicted interactions (Fig. 6A) revealed an interesting link between early and late GRNs. Early-expressed repressor TFs like *Nfib* and *Bach2* (P5, see Fig. 2D) target later TFs (P15-28), such as *Bcl6* and *Smad3* (Fig. 6A). This suggests that early GRNs could repress later ones, potentially delaying the activation of their downstream targets. Once active, late GRNs are also predicted to target each other, as seen with *Bcl6*, *Klf9* and *Foxo1* targeting *Bcl11b*, or *Klf9* regulating *Bcl11b*, *Nr3c1* and *Smad3* (Fig. 6A, Fig. S7). Additionally, early-expressed activator TFs like *Rfx3* are predicted to target later TFs, such as *Bcl6*, and in some cases these predictions are bidirectional, suggesting feedback between early and late GRNs that would mutually regulate each other, e.g. Rfx3(+)-Klf9(+)-Bcl11b(+) (Fig. 6A). To link the GRNs to their putative roles in synapse development, we used SynGO to determine the size of each node based on the percentage of eRegulon targets present in the database. Late-active eRegulons (clusters 2 and 3) show a higher proportion of SynGO-annotated synaptic genes than early-active eRegulons (cluster 1) (Fig. 6A, Fig. S7), with Atf6(+) having the highest (40%) and Rfx3(-) the lowest (4.7%) proportion.

**Fig 6.**
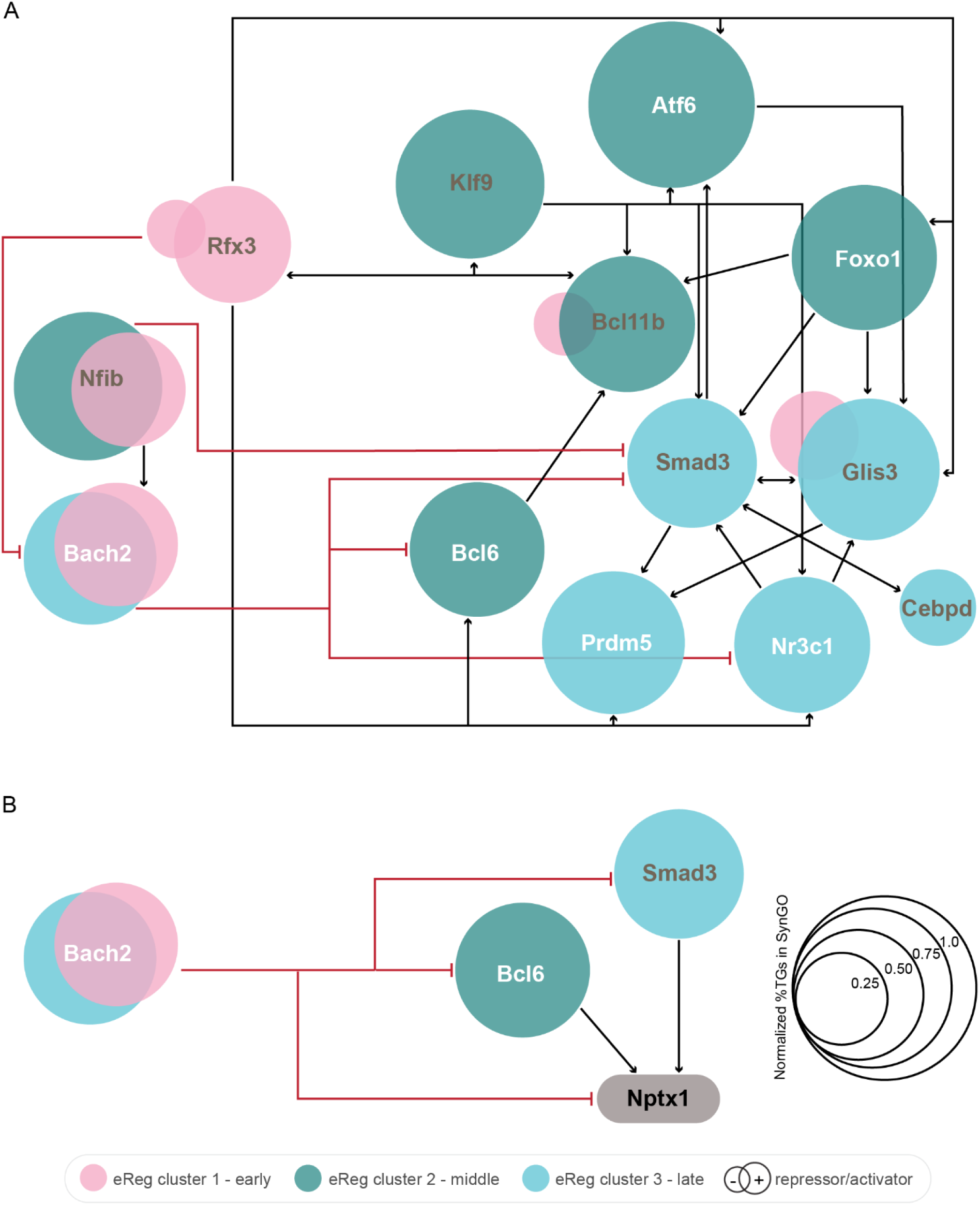
Schematic representation of gene regulatory network (GRN) model centered on Bcl6(+) and Smad3(+) eRegulons. **A)** Simplified GRN model (see also Fig. S7) centered on Bcl6(+) and Smad3(+) eRegulons and the predicted interactions of leading transcription factors (TFs) (incoming and outgoing; activating (black lines) and repressing (red lines)). eRegulons are represented by circles, colored depending on the cluster the eRegulon belongs to (cluster 1-early, pink; cluster 2-middle, green; cluster 3-late, blue; see also Fig. 2D, Fig. S3G). Early-expressed repressor TFs *Nfib* and *Bach2* target later TFs *Bcl6* and *Smad3*. **B)** Simplified representation of *Bach2*, *Bcl6* and *Smad3* interactions with their target gene (TG) *Nptx1*. In both A and B, the size of the circles for each eRegulon is determined based on the percentage of SynGO-annotated synaptic TGs, normalized to the eRegulon with the highest proportion of SynGO-annotated synaptic genes, Atf6(+). eRegulons with leading TFs that act as both activators and repressors are represented by two circles, with the left circle representing the repressor, and the right circle representing the activator. TFs labeled in grey also target themselves (e.g. *Nfib*), while those labeled in white do not (e.g. *Bach2*).

Early repression of late-expressed genes was also observed beyond TFs. For instance, the synaptic organizer-encoding gene *Nptx1*, a shared predicted target in Bcl6(+) and Smad3(+) eRegulons (see Fig. 4C-D and Fig. 5B), is also a predicted target of the Bach2(-) repressor (Fig. 6B). Similarly, *Tanc1*, a Smad3(+) target involved in synaptic organization (*65*) (Fig. 5B), is predicted to be repressed by Nfib(-).

These observations suggest a key role for early TFs in repressing late TFs and their target genes controlling synaptic organization and maturation. Cross-referencing the identified TFs with SFARI (*68*), a curated database of Autism Spectrum Disorder (ASD) genes, showed that approximately 80% of the ASD-associated TFs in our dataset belong to the early cluster of GRNs (cluster 1; including *Mef2c*, *Nfi* family members, *Zbtb20* and *Rfx3* (Fig. S8, Table S1)). In contrast, no ASD-associated TFs were identified in the late cluster (Fig. S8, Table S1). This suggests that the early TFs regulating GC synapse development are susceptible in ASD. Together, these findings highlight a sequential regulatory program in which early-active TFs act as gatekeepers, delaying the activation of later GRNs and their putative synaptic targets.

## DISCUSSION

Here, we combine transcriptomics, epigenomics, and translatomics to reconstruct the GRNs regulating synaptic development of GCs. Our multiomic analysis reveals distinct GRNs active at different stages of GC development and identify the TFs *Bcl6* and *Smad3* as regulators of GC synapse development. In addition, our results suggest that early-active TFs repress GRNs that are active at later stages of GC development.

We find a dynamic gene regulatory code orchestrating postnatal development of GCs as they integrate into the hippocampal circuit. GCs undergo prominent shifts in gene expression between early (P5-P7) and late (P15-P28) timepoints at both the transcriptomic and translatomic levels, with most genes upregulated in the early phases and downregulated in the late phases. Genes upregulated in early stages are key for neurite development, while genes active at later timepoints control synaptic processes. GRN inference in mGCs identifies 36 eRegulons, which can be clustered into three main temporal groups (early (P5), middle (P10-15) and late (P15-28)). Among the TFs with established roles in the DG, we find *Bcl11b*, *Mef2c* and *Klf9*, all three of which are implicated in the development of GC connectivity (see refs (*28*, *31*, *32*)). Half of the TFs leading the eRegulons we identify have no known role in the DG (Table S1).

We observe interesting parallels in GRN structure of postnatally developing GCs (this study) and adult-born GCs (aGCs) (*47*). In the adult DG, radial-glia like cells continuously generate new neurons, which functionally integrate into the hippocampal circuit (*69*, *70*). These aGCs are more excitable but less precisely-tuned than their embryonic-born counter parts (*71*). SCENIC (*52*) analysis of scRNAseq data from immature and mature aGCs identified 9 regulons (*47*) that overlap with our findings, including Bcl6(+), and Foxo1(+) (Table S1). Conversely, some eRegulons identified in our study are led by TFs previously reported to play a role in adult neurogenesis, such as *Mef2a* (*72*), *Neurod2* (*73*), and *Foxo1* (*74*). Together, these results suggest shared gene regulatory mechanisms between perinatal and adult-born GC development.

In our functional analysis of the role of identified GRNs in synaptic development of GCs, we characterize the *Bcl6* TF as a novel regulator of structural maturation of GC synapses. Previous work in the mouse cortex showed that *Bcl6* promotes neurogenesis via repression of Notch targets (*57*), but a role in postnatal synapse development has not been described. Here, we identify *Bcl6* as a transcriptional activator targeting 23 genes that show similar expression patterns as *Bcl6*, further supporting an activator role for *Bcl6* in postnatal GCs. TFs can have dual repressor/activator roles, such as *Fezf2* in corticospinal neurons (*20*), including TFs exerting their transcriptional regulation via ZiN/POZ (poxvirus zinc finger)-domains like *Bcl6* (*75*). We found that *Bcl6* KD reduces density of mature dendritic spines in GCs. Several of the Blc6(+) predicted target genes regulate dendritic spines. KD of *Pacsin1* in CA1 neurons reduces spine density (*59*), while overexpression of *Arhgef25* increases spine size and the proportion of mature spines (*60*). Our observations suggest that *Bcl6* is necessary for the maturation of dendritic spines in GCs, as mushroom spines are considered the mature type associated with stronger synapses critical for memory storage (*76*). In addition to the effects on dendritic spine density, *Bcl6* KD increases filopodia density in MFBs, while other MFB features such as area and volume remain unaffected. MFB filopodial synapses target GABAergic cells in the stratum lucidum of CA3 (*64*), mediating feedforward inhibition essential for CA3 microcircuit function in normal and pathological conditions (*8*, *77*, *78*). MFB filopodia density peaks during development (P14) and decreases in adult mice (*39*), suggesting that loss of *Bcl6* impairs maturation or refinement of filopodial density. Several synaptic genes affect MFB filopodia number (*8*, *9*, *78*), and some of the Bcl6(+) predicted target genes could influence filopodial growth, such as the actin cytoskeleton organizer *Pkp4* (*79*). Together, these findings suggest that *Bcl6* promotes structural maturation of GC synapses through its interactions with specific synaptic target genes.

In contrast, loss of *Smad3* in GCs does not affect dendritic spine density or MFB morphology. These results were surprising, considering that the Smad3(+) eRegulon contains 224 predicted target genes with roles in synapse organization and plasticity (*10*, *61*, *65*). Further analysis revealed a selective increase in mIPSC frequency in *Smad3* KO GCs. This increase could reflect an increase in density of inhibitory input onto GCs or a change in inhibitory presynaptic properties. GCs in adult *Smad3* KO mice show increased inhibitory, but not excitatory, neurotransmission (*54*), consistent with our analysis of cell-autonomous loss of *Smad3* in developing GCs. Taken together, our results suggest that the Bcl6(+) and Smad3(+) eRegulons control distinct aspects of GC synapse development.

Our analysis of interactions between GRNs predicts sequential regulations, with early-active TFs delaying the activation of later GRNs and their putative synaptic targets. This regulation is exemplified by the interactions of the Bcl6(+) and Smad3(+) eRegulons. Both eRegulons are targeted by early-active repressor GRNs, such as Nfib(-) and Bach2(-). *Nfib* loss in the cortex induces severe loss of axonal projections and upregulation of Notch signaling genes (*80*), suggesting a repressor role for this TF. Our GRN model (Fig. 6, Fig. S7) suggests that early-active Nfib(-) can repress Smad3(+), through shared target genes such as *Tanc1*. The *Bcl6*-*Bach2* link has been previously studied in immune cells. *Bcl6* is repressed by *Bach2* in T-cells (*81*), while *Bcl6* and *Bach2* cooperate to repress other genes in B-cells (*82*). *Bach2* controls the selection of viable cells via the activation of the tumor suppressor protein p53 in the absence of *Bcl6*, and their dysregulation is associated with high leukemic cell proliferation (*83*, *84*). A similar antagonistic relationship between *Bach2* and *Bcl6* may regulate GC synapse development. Our GRN model (Fig. 6, Fig. S7) suggests that early-active Bach2(-) can repress the Bcl6(+) and Smad3(+) eRegulons through shared target genes such as *Nptx1* (Fig. 6B).

The terminal selector hypothesis proposed in *Drosophila* and *C. elegans* studies suggests that combinations of constantly expressed TFs define key cellular properties such as neurotransmitter identity and connectivity (*15*, *18*, *85*, *86*). The GRNs belonging to the middle and late clusters we identify here differ from terminal selectors in their postnatal onset of activity, after the wiring pattern of GCs is established. Cell type-specific combinations of such GRNs could enable neurons to develop a large synaptic diversity without altering fundamental properties such as neurotransmitter profiles or wiring patterns. Strikingly, we find that ±80% of the early-active TFs we identify are associated with ASD, suggesting that the early GRNs regulating GC development are particularly susceptible in ASD. Whether this observation extends to other hippocampal neuron types remains to be determined.

In conclusion, our results uncover a dynamic gene regulatory code governing synaptic development of GCs, provide insight in key TFs and target genes, and identify *Bcl6* and *Smad3* as regulators of specific aspects of GC synapse development. Our single-cell transcriptome, single-nucleus multiome, and GC translatome datasets provide an invaluable resource for the poorly mapped postnatal developmental period of hippocampal GCs. A systematic extension of our multiomic approach to other cell types during postnatal development will be critical to uncover the gene regulatory codes driving synapse formation, maturation and plasticity in brain regions such as hippocampus and cortex.

## MATERIALS AND METHODS

### Animals

All animal experiments were conducted according to the KU Leuven ethical guidelines and approved by the KU Leuven Ethical Committee for Animal Experimentation. Mice were maintained in a specific pathogen-free facility under standard housing conditions with continuous access to food and water. Wild-type (WT) C57BL/6J mice originated from Charles River. Rpb4-Cre (MGI:4367067) and Ai9 (JAX:007909) mouse strains were obtained from Dr. Aya Takeoka (NERF, Leuven). RiboTag (JAX:029977) and H11-Cas9 line (JAX: 028239) were obtained from JAX. Rosa26-LSL-Cas9 (JAX:024857) mouse line was obtained from Dr. Jan Cools (VIB Center for Cancer Biology (CCB), Leuven). Genotypes were regularly checked by PCR analysis. Rbp4-Cre heterozygous Cre positive males were bred to WT, Ai9, RiboTag or Rosa26-LSL-Cas9 females. For euthanasia, animals were either anesthetized with isoflurane, and decapitated or injected with an irreversible dose of ketamine–xylazine.

### Single-cell RNA seq

(scRNAseq) Rbp4-Cre:Ai9 mice were sacrificed at different developmental stages (postnatal (P) day 5, P7, P10, P15, P28), after being anesthetized with isoflurane. Neuronal extraction protocol was adapted from Allen Brain Neuronal Isolation Protocol (*87*), and Tasic et al, Nat Neuroscience, 2016 (*88*). Briefly, immediately after extraction the brain was immersed in ice cold carbogenated artificial cerebrospinal fluid (ACSF) (Table S2). The brain was sliced in a pre-cooled brain matrix (AgnThos 69-2165-1) to 500 μm sections, and the DG was dissected in a petri dish containing ice-cold ACSF while carbogenating. Immediately after dissection the DG pieces were transferred into an enzymatic solution containing Pronase Protease (VWR 537088, 2 mg/ml), diluted in digestion medium (DM) (Table S2), and supplemented with 50 mM Trehalose (Sigma T0167). The enzymatic solution was placed in a 37 °C water bath while carbogenating, incubated for: 10 minutes (min) (for P5, P7 and P10), 15 min (for P15), 28 min (for P28) followed by the addition of Trypsin (Sigma 59417C, 0.125%) and DNaseI (Fisher Scientific 10649890, 0.2 mg/ml) for 2 min (except for the P28 sample, which received no trypsin nor DNaseI). After digestion, the enzymatic solution was removed, and tissue pieces were washed with 10 ml DM (ice-cold and carbogenated). The tissue was triturated using Pasteur pipettes in 2ml of Opti-MEM/trehalose solution (Table S2) with polished tip openings of 500-μm and 250-μm diameter, layered on top of a 20% Percoll solution (Table S2) and spun at 700g, for 10 min at 4 °C. When all solution and debris was removed, the cell pellet was resuspended in 200-500 μl Opti-MEM/trehalose solution (Table S2).

Single cells were isolated by FACS into individual wells of 96-well plates (4titude, Cat. No. 4TI-0960/C) containing 4 μl of cell lysis buffer (0.1% Triton X-100, Sigma Aldrich; 1U/μl of RNase Inhibitor, RNase OUT, Thermo Fisher; 2.5 μM Oligo dT (25), IDT; and 2.5 mM dNTP, Promega). Sorting was performed on a BD FACSAria Fusion using a 100 μm nozzle, a sheath pressure of 20 psi, and in the single-cell sorting mode. To exclude dead cells, DAPI (Invitrogen 10184322) was added to the single cell suspension to a final concentration of 5 ng/ml. FACS populations were chosen to select cells with low DAPI and high tdTomato fluorescence. The cells were sorted into wells with lysis buffer and the plate was spun down at 2000xg for 1 min prior to storage at -80 °C.

### Library preparations and sequencing for single-cell RNA-seq

Smart-seq2 was performed using a previously published protocol (*45*). For the first strand synthesis, the Smart-Seq2 plates were thawed to room temperature for 1 min and subsequently spun down at 2000 xg for 1 min. The RNA denaturation was carried out at 72 °C for 10 min and flash-cooled on ice for 5 min. First strand synthesis mix (1X SuperScript II first strand synthesis buffer, Thermo Fisher; 5 mM DTT, Thermo Fisher; 100 U Oligo SuperScript II reverse transcriptase enzyme, Thermo Fisher; 10 U RNAse OUT, Thermo Fisher; 1M Betaine, Sigma Aldrich; 6 mM MgCl2, Thermo Fisher; and 1 μM TSO, IDT Technologies) is added to the cell lysis mix to a total volume of 10 μM. First strand synthesis reaction was carried out with following program: 42 °C for 90 min; 10 cycles of {50 °C for 2 min; 42 °C for 2 min}; 72 °C for 15 min; 4 °C indefinite hold. PCR amplification of the first strand product was performed by adding 15 µl of the PCR mix to the first strand product (1X KAPA HiFi HotStart Ready Mix, Roche; and 0.1 µM of IS PCR primer, IDT Technologies). The 22 cycles of the following PCR program was used to amplify the first strand product: 98 °C for 3 min; 22 cycles of {98 °C for 20 s; 67 °C for 15 s; 72 °C for 6 min}; 72 °C for 5 min; 4 °C indefinite hold. Using Hamilton STAR liquid handler (Hamilton), 20 µl of Ampure XP was added to each well and mixed. Standard Ampure XP purification was carried out as per the manufacturer’s recommendation and the cDNA library was eluted in 22.5 µl of Elution buffer. 22 µl of the eluted library was transferred to fresh Echo Mass Spectrometry (MS) Qualified 384-well Polypropylene (PP) Microplate (Beckman Coulter). Labcyte Echo 550 acoustic dispenser (Beckman Coulter) was used for the next steps. 1 µl of Tn5 tagmentation mix (1X Tagment DNA buffer, Illumina; ATM mix, Illumina and cDNA library 1 ng) for cDNA fragmentation. Tn5 tagmentation was performed with the following program: 55 °C for 10 min; 4 °C indefinite hold. Tagmentation reaction was stopped by quenching the reaction with 500 nl of NT buffer for 5 min (Illumina). 750 nl of the i5 and i7 Illumina indexes were added to the plates. Finally, 2 µl of NPM master mix was added to the plate and mixed well and place for index PCR amplification: 72 °C for 3 min; 95°C for 30 s; 12 cycles of {95 °C for 10 s; 55 °C for 30 s; 72 °C for 30 s}; 72 °C for 5 min, and 4 °C indefinite hold.

The indexed PCR products are pooled in a single tube and 0.8X Ampure XP purification (Beckman Coulter) was carried out as per manufacturer’s recommendation and finally the sequencing library was eluted in 35.5 µl of Elution buffer (Qiagen). Smart-Seq2 libraries were sequenced on NextSeq 500 or NextSeq2000 (Illumina) sequencing platforms. Sequencing was done as per the protocol recommendations: paired end read of 61 bps (read 1), 61 bps (read 2), 8 bps (index 1) and 8 bps (index 2). Sequencing was performed at a depth of 500,000 reads per cell.

### Nuclei isolation and library preparation for 10X Genomics single-cell multiome

Nuclei preparation from mouse hippocampal tissue was done using sucrose gradient protocol. Briefly, up to 50 mg of hippocampal tissue was homogenized using 1 ml of Homogenization Lysis Buffer (HLB) (1X salt-Tris buffer: 146 mM NaCl, 10 mM Tris 7.5, 1 mM CaCl_2_, 21 mM MgCl_2;_ 0.03 % v/v Tween-20, Sigma Aldrich; 0.01% v/v NP40, Sigma Aldrich; 1% BSA, Thermo Fisher; 25 mM KCl, Thermo Fisher; 250 mM Sucrose, Sigma Aldrich; 1mM DTT, Thermo Fisher; 1X Protease Inhibitor, Sigma Aldrich; 1 U/ul RNAse Inhibitor, Promega). The tissue was homogenized with 10X strokes with pestle A and 10X strokes with pestle B. The homogenized mixture was incubated in HLB for 5 min. During the incubation, the homogenized sample was passed through 70 µm nylon mesh. The nuclei were pelleted by centrifugation at 500xg for 5 min. The nuclei pellet was washed and resuspended with 520 µl of Wash Buffer 1 (1X salt-Tris buffer; 1% BSA, Thermo Fisher; 25 mM KCl, Thermo Fisher; 250 mM Sucrose, Sigma Aldrich; 1mM DTT, Thermo Fisher; 1X Protease Inhibitor, Sigma Aldrich; 1 U/µl RNAse Inhibitor, Promega). The nuclei suspension was then mixed with equal volume of Gradient Medium (1 mM CaCl_2_, Thermo Fisher; Optiprep 50% v/v, Sigma Aldrich; 5 mM MgCl_2_, Thermo Fisher; 10 mM Tris 7.5, ThermoFisher; 75 mM Sucrose, Sigma Aldrich; 1mM DTT, Thermo Fisher; 1X Protease Inhibitor, Sigma Aldrich; 0.5 U/µl RNAse Inhibitor, Promega). This was thoroughly mixed, and the mix was layered upon 948 µl of 29% density cushion (Optiprep 29% v/v, Sigma Aldrich; 72.5 mM KCl, Thermo Fisher; 14.5 mM MgCl_2_, Thermo Fisher; 29 mM Tris 8, Thermo Fisher; 121 mM Sucrose, Sigma Aldrich; 0.25 U/µl RNAse Inhibitor, Promega). The layered nuclei suspension was subjected to centrifugation at 9000xg for 20 min at 4 °C. After the centrifugation, the debris was removed from the top of the suspension and if excess debris was present, this could be scooped out with a 1000 µl pipette tip. The supernatant was aspirated out gradually using a glass Pasteur pipette until about 75 µl of the supernatant was left. Equal volume of 0.2X Lysis Buffer (20 mM Tris 7.5, ThermoFisher; 20 mM NaCl, ThermoFisher; 6mM MgCl_2_, ThermoFisher; 0.02% Tween 20, Sigma Aldrich; 0.02% NP40, Sigma Aldrich; 0.002% Digitonin, Promega; 2% BSA, Thermo Fisher; 2mM DTT, ThermoFisher; 2U/µl Protector RNAse Inhibitor, Sigma Aldrich) was added and the nuclei were resuspended and incubated for 2 min on wet ice. After the incubation the lysis was quenched by adding 1 ml of Wash Buffer 2 (10 mM Tris 7.5, ThermoFisher; 10 mM NaCl, ThermoFisher; 3 mM MgCl_2_, ThermoFisher; 0.01% Tween 20, Sigma Aldrich; 1% BSA, Thermo Fisher; 1 mM DTT, ThermoFisher; 1 U/µl Protector RNAse Inhibitor, Sigma Aldrich) and the nuclei suspension was resuspended. The nuclei were pelleted by centrifugation at 500xg for 5 min. The supernatant was discarded until about 20 µl of the supernatant was left in the Eppendorf. Removal was done carefully without disturbing the nuclei pellet. Nuclei pellet was resuspended with 20 µl of 2X Nuclei Buffer (2X Nuclei buffer, 10X Genomics, PN-2000153; 1 mM DTT, ThermoFisher; 1U/µl Protector RNAse Inhibitor, Sigma Aldrich).

The nuclei count and the viability of the samples were accessed using LUNA dual florescence cell counter (Logos Biosystems) and a targeted cell recovery of 10000 cells was aimed for each of the samples. Library preparations for 10X Genomics single cell Multiome were performed using 10X Genomics Chromium NextGEM Single cell Multiome ATAC+Gene Expression chemistry (10X Genomics, Pleasanton, CA, USA). Transposition of nuclei and the rest of the 10X Genomics Multiome library preparation was performed as per the protocol recommendations (Chromium Next GEM Single Cell Multiome ATAC + Gene Expression Reagent Kits User Guide; CG000338 Rev G). At the different check points the library quality was accessed using Qubit (ThermoFisher) and Bioanalyzer (Agilent).

10X libraries were sequenced either on Illumina platform or on MGISEQ-2000 sequencing platform (MGI Hong Kong SAR sequencing facility). Before being able to sequence on the MGI platforms, the 10X libraries need to undergo a conversion step using MGIEasy Universal DNA Library Prep kit. Briefly, the final 10X sequencing libraires were circularized using the splint-ligation step and the circularized libraries were converted to single-stranded DNA copies. DNA nanoballs were prepared from the circularized ssDNA using Rolling Circle Amplification (RCA). DNB libraries generated were then flown through the patterned flow cell of the MGISEQ-2000RS High-Throughput Sequencing kit. For a targeted sequencing saturation of 50-60%, sequencing was performed at a depth of 30,000 - 60,000 reads per cell for the RNA and ATAC libraries. For RNA libraries sequencing recipe of paired end read of 28 bps (read 1), 100 bps (read 2) and 10 bps (index 1) and 10 bps (index 2) was used for sequencing. For ATAC libraries sequencing recipe of paired end read of 50 bps (read 1), 49 bps (read 2) and 24 bps (index i5) and 8 bps (index i7) were used for sequencing.

### RNAscope in situ hybridization

Rbp4-Cre:Ai9 mice were sacrificed at different developmental stages (P5, P15, P28), brains were dissected and frozen in OCT (4583, Sakura Finetek). 10 µm cryosections were prepared on a CM3050 S cryostat (Leica), mounted on SuperFrost Ultra Plus adhesion slides (Thermo-Fisher) and dried at RT. Sections were stained using the RNAscope™ Multiplex Fluorescent Detection Kit v2 (ACDbio, 323110) following manufacturer protocols. Briefly, sections were treated for 10 min with hydrogen peroxide solution and 12 min with protease IV with 1X PBS washing in between. RNAscope probes were hybridized at 40 °C for 2h and subsequently HRP signal developed using OPAL dyes. *Bcl6* probe was used in the C1-channel (+ Opal 520 1:2000), *Smad3* probe was used in the C2-channel (+ Opal 520 1:2000), while a *tdTomato* probe was used in the C3 channel (+ Opal 570 1:2000) (Table S3). Sections were counterstained with DAPI for 30 seconds (s) and mounted with Mowiol. Images of *tdTomato* positive regions in the DG were acquired on a Zeiss LSM900 using the 20X objective. Constant laser power and gain settings were used to image the same probe in sections of the three different ages. Images were quantified by measuring probe mean intensity over area of a *tdTomato*-positive ROI using ImageJ. Statistical analyses were performed using GraphPad Prism 9.

### RiboTag sample preparation and bulk RNA sequencing

Rbp4-Cre:RiboTag Cre mice were sacrificed at different developmental stages (P5, P7, P10, P15, P21, P28) and hippocampi were dissected in ice-cold HBSS and frozen for subsequent use. Hippocampi from male and female mice were used at all timepoints, 3-4 animals were pooled per sample at P5 and P7, and 2 animals per sample were pooled at P10, P15, P21 and P28. Moreover, control samples from P10 animals negative for Cre-recombinase were used to ensure specificity of the pull-down.

RiboTag sample preparation protocol was adapted from Sanz et al, 2019 (*89*). Briefly, hippocampi were thawed in pre-chilled homogenization buffer containing 50 mM TrisCl pH 7.5, 100 mM KCl, 12 mM MgCl_2_, 1% NP40, 1 mM DTT, 200 U/ml RiboLock RNase Inhibitor (Thermofisher, EO0382), 1 mg/ml Heparin (Sigma, H3393), 0.1 mg/ml cycloheximide (Sigma, C4859), 1X EDTA-free protease inhibitors. Next, samples were placed in a Dounce homogenizer where they were dissociated using 2 sizes glass pestles. Lysate was centrifuged at 10000g, 4 °C for 10 min and supernatant (named hereafter as input) was kept for further processing. 40 μl Pierce™ Anti-HA Magnetic Beads (ThermoFisher, 13464229) were added to the input and incubated overnight at 4 °C on a microtube rotator with gentle mixing. Next, samples were placed on magnetic stand for at least 1 min to remove supernatant (named hereafter as unbound), and washed 4x using high salt buffer (50 mM TrisCl pH 7.5, 300 mM KCl, 12 mM MgCl2, 1% NP40, 0.5 mM DTT, 0.1 mg/ml cycloheximide). After the last wash, 350 μl RLT buffer (Qiagen RNeasy mini kit, 74104) supplemented with β-mercaptoethanol was used to elute RNA from the ribosomes. The mix was vortexed 30s and tubes were placed back in magnetic stand to collect the clear RLT buffer containing the RNA. Next, RNA was isolated using Qiagen RNeasy mini kit with DNase treatment and following manufacturer instructions.

### RiboTag quality control: bioanalyser, quantitative RT-PCR and western blot

*Bioanalyser* RNA quality and concentration was checked on a Bioanalyser instrument (Agilent Technologies) using High sensitivity RNA ScreenTape (Agilent, 5067-5579). Only RNA samples with RNA integrity number (RIN) value higher than 7.5 were used.

*Quantitative RT-PCR* For quality control using qPCR, 60-120 ng RNA (P5, P10, P15, n=2 samples per condition) were retrotranscribed using High-Capacity cDNA Reverse Transcription Kit (Thermofisher, 4368814). To determine the fold-enrichment of respective marker genes in immunoprecipitated RNA compared with P10 whole hippocampal RNA lysates, LightCycler® 480 SYBR Green I Master kit (Roche) and comparative C^T^ method were used. Samples were considered specific if immunoprecipitated RNA exhibited correct de-enrichments or enrichments of respective marker genes and if RNA of control samples did not show any selectivity for marker genes (see Table S4 for primer sequences). For each assay, three technical replicates were performed, and the mean was calculated. The mRNA levels were normalized to *Pgk1* mRNA (housekeeping gene) (Table S4). Quantitative RT–PCR assays were analyzed using GraphPad Prism 9.

*Western blot* Input fraction of hippocampal samples were collected for n=2 P5, P10, P15 Rbp4-Cre:RiboTag and n=2 P10 RiboTag mice (labeled as CTRL, Cre neg). 2% volume of input was used for western blot. Briefly, samples were boiled for 5 min at 95 °C, samples were loaded on a 10% pre-cast polyacrylamide gel (Bio-Rad, 4561036) and run at 180V. Proteins were transferred to 0.2 μm nitrocellulose membrane using semi-dry transfer (Biorad) using the mixed-molecular weight program. Membranes were blocked in 5% milk in TBS-T buffer (25 mM Tris-base pH 7.5, 300 mM NaCl, 0.05% Tween-20) for 1h at RT. Primary and secondary antibodies were diluted in 5% milk in TBS-T. Primary antibodies were incubated O/N at 4 °C, secondary antibodies were incubated 2h at RT (Table S3). After primary and secondary antibody incubation, membranes were washed 5 times with TBS-T and once with 1X PBS before imaging on ImageQuant 800 (Cytiva) using SuperSignal West Femto PLUS Chemiluminescent Substrate (Thermo Fisher 34577).

### RiboTag library preparation and sequencing

Library preparation from total RNA extraction and sequencing was performed at the VIB Nucleomics Core. Per sample, an amount of 25-100 ng of total RNA was used as input. Using the Illumina TruSeq® Stranded mRNA Sample Prep Kit (protocol version: #1000000040498 v00 October 2017) poly-A containing mRNA molecules were purified from the total RNA input using poly-T oligo-attached magnetic beads. In a reverse transcription reaction using random primers, RNA was converted into first strand cDNA and subsequently converted into double-stranded cDNA in a second strand cDNA synthesis reaction using DNA Polymerase I and RNAse H. The cDNA fragments were extended with a single ’A’ base to the 3’ ends of the blunt-ended cDNA fragments after which multiple indexing adapters were ligated introducing different barcodes for each sample. Finally, enrichment PCR was carried out to enrich those DNA fragments that have adapter molecules on both ends and to amplify the amount of DNA in the library. Sequence libraries of each sample were equimolarly pooled and sequenced on Illumina NovaSeq 6000 (v1.5, SE100 (100-8-8-0), 1% PhiX).

### Cell Lines

HEK293T-17 human embryonic kidney cells were obtained from American Type Culture Collection (ATCC) cat# CRL-11268. HEK293T-17 cells were grown in Dulbecco’s modified Eagle’s medium (DMEM; Invitrogen) supplemented with 10% fetal bovine serum (FBS; Invitrogen) and penicillin/streptomycin (Invitrogen).

### Plasmids

shRNA sequences for *Bcl6* knockdown (KD) and control were obtained from Tiberi et al, 2012 (*57*) (Table S5). Hairpin sequences (fwd tcaagagagattctcagatccgtgtc, rev ctcttgaagattctcagatccgtgtc) were obtained from Karaca et al, 2018 (*90*). gRNA sequences for *Smad3* knockout (KO) were generated using CRISPick database with the Mouse GRCm38 (NCBI RefSeq) chosen as reference genome, CRISPRko as mechanism and the SpyoCas9 (Hsu 2013) tracrRNA as enzyme. Two gRNA sequences with a Target Cut% below 30% were selected per target gene (Table S5). For both gRNA and shRNA forward and reverse oligos were designed and SapI (NEB) restriction site overhangs were added.

For the Supernova sparse labeling, pAAV-TRE-DIO-FLP (Addgene 118027) and pAAV-TRE-FLP (Lab of Synapse Biology, from Addgene 118027) were used for *Smad3* KO and *Bcl6* KD, respectively. shRNAs were cloned into the pAAV-U6-TRE-fDIO-MARCKS-EGFP-IRES-tTA plasmid (Lab of Synapse Biology, from Addgene 118026). mGFP is tagged with a MARCKS (myristoylated alanine-rich protein kinase C substrate) sequence to be targeted to the membrane. gRNAs were cloned into the pAAV-U6-TRE-fDIO-MARCKS-mScarlet3-IRES-tTA plasmid (Lab of Synapse Biology, from Addgene 118026 with mScarlet3 sequence from Addgene 189753). pRep/Cap (AAV2/9, Addgene 112865) and pΔF6 (Addgene 112867) plasmids were used to generate AAVs.

### AAV production

High-titer AAV production and purification was carried out as previously reported (*91*). Briefly, 6 plates of 80% confluent HEK cells were transfected with 20 µg of pDelta F6 plasmid, 10 μg of pRepCap 2/9 plasmid and 10 μg AAV genome plasmid per plate using PEI transfection. 3 days later cells were collected by scraping and centrifugation. AAV particles in the supernatant were precipitated by PEG and added to cell lysates. Lysates were treated by Benzonase nuclease to remove cellular DNA. Lysates were loaded onto an iodixanol gradient (60%, 40%, 25%, 15%) and ultracentrifuged 1:40 hours at 50.000 RPM at 12 °C. The 40% fraction containing the purified AAV was carefully collected and desalted and concentrated on a 4 ml Amicon column (Sigma). AAV purity was tested by silver staining using the ProteoSilver™ Silver Stain Kit (Sigma PROTSIL1-1KT). AAV titer was determined by qPCR to normalize titers between different viral vector batches using LightCycler 480 SYBR Green I Master (Roche 04707516001). Primers for qPCR targeting the synapsin promoter were as follows: fwd TGATAGGGGATGCGCAATTTGG, and rev GTGCAAGTGGGTTTTAGGACCA. Each AAV solution for KO/KD combined two different viruses, each carrying a different gRNA or shRNA, to enhance efficiency.

### Stereotactic injections

For *Smad3* loss-of-function analysis Rbp4-Cre:Rosa26-LSL-Cas9 mice littermates were injected at P7. P7 mice were anesthetized with 5% isoflurane and placed in a mouse stereotact (KOPF) on a hot plate kept at 37 °C. During the rest of the procedure 2% isofluorane was constantly administered. After disinfecting the mouse’s head, local anesthesia was administered by a subcutaneous injection with 100 μl lidocaine (xylocaine 1%). An incision was made on the skin to reveal the skull. One bilateral injection was performed, coordinates to target DG at P7 were as follows: x: ±1.3, y: 1.4 from bregma, z: -2.1 from the surface of the skull. 100 nl of AAV mix containing AAV-TRE-DIO-FLP:AAV-gRNA-mScarlet in a 200:1 ratio were injected using a Nanoject III (Drummond) through a beveled capillary at 5 nl/s. After a 2 min recovery, the capillary was pulled out at ∼0.1 mm/5 s. The incision was stitched with surgical glue (Millpledge Veterinary). After 6 h, their health was examined, and mice were injected with 0.1 mg/kg buprenorphine.

For *Bcl6* loss-of-function analysis C57Bl6J littermates were injected at P0. P0 mice were anesthetized using hypothermia-induced anesthesia for 15 min and kept on an ice-cold surface during the procedure. The pups were immobilized manually and 50 nl of AAV mix containing AAV-TRE-FLP:AAV-shRNA-mGFP in a 100:1 ratio was injected using a Nanoject III (Drummond) through a beveled capillary at 5 nl/s. After 30 s recovery, the capillary was pulled out at ∼0.1 mm/5s. Two injections per hemisphere were performed to target DG at P0, the x and y coordinates were visually determined and a z: -1.6 from the surface of the skull was used.

### Tissue processing and immunofluorescence

Mice were trans-cardially perfused with 10 ml of 4% sucrose 2% sucrose in 1X phosphate buffer (0.1 M, pH 7.4) under non-reversible anesthesia. Brains were dissected and post-fixed for 1h in PFA, washed in PB and sliced to 80 μm thick by vibratome. Sections were blocked in 10% normal horse serum (NHS), 0.5M Glycine, 0.5% Triton-X 100 in PBS before incubation with antibodies in 5% NHS, 0.5% Triton-X 100 in PBS with extensive washing between primary and secondary antibodies (Table S3). Sections were counterstained with DAPI before mounting on microscope slides with Mowiol. Imaging was performed on a Zeiss LSM900 using airyscan mode.

### Neuronal cultures

Hippocampal or cortical cultures were obtained from dissected E18 H11-Cas9 mice or C57Bl6J WT embryos and plated on poly-D-lysine (Millipore) and laminin (Invitrogen)-coated 6-well plates. Neurons were maintained in Neurobasal medium (Invitrogen) supplemented with B27. 48-72 h after seeding, 10 mM glia-inhibitor (5-Fluoro-2-deoxyuridine; Sigma-Aldrich) was added to the medium. To achieve *Bcl6* KD, WT neurons were infected at DIV1 with high-titer AAVs expressing either shRNAs against the target gene or control shRNAs and collected at DIV15 for qPCR. To achieve *Smad3* KO, H11-Cas9 neurons were infected at DIV1 with high-titer AAVs expressing either gRNAs against the target gene or control gRNA against lacZ and protein was collected at DIV15 for western blot.

### Validation of shRNAs and CRISPR gRNAs (WB and RT-PCR)

*CRISPR KO validation* Primary neuronal cultures from H11-Cas9 mice were infected at DIV1 with 1 μl of a mix of AAV-U6-*Smad3*-gRNA-TRE-fDIO-MARCKS-mScarlet3-IRES-tTA (gRNA1 + gRNA2) (1:2000) or 1 μl AAV-U6-lacZ-gRNA-TRE-fDIO-MARCKS-mScarlet3-IRES-tTA (1:2000). Neurons were lysed at DIV15 using RIPA buffer (50 mM Tris-HCl, 150 mM NaCl, 1% Triton-X 100, 0.5% sodium deoxycholate, 1% SDS). 50-100 ng of protein lysate were loaded on a 10% pre-cast polyacrylamide gel to evaluate KO efficiency between conditions (see Table S4 for antibodies).

*shRNA KD validation* Neuronal primary cultures from C57Bl6J mice were infected at DIV1 with 1 μl of a mix of AAV-U6-*Bcl6*-shRNA-TRE-fDIO-MARCKS-EGFP-IRES-tTA (shRNA1 + shRNA2) (1:2000) or 1 μl AAV-U6-scrambled-shRNA-TRE-fDIO-MARCKS-EGFP-IRES-tTA (1:2000). Neurons were lysed at DIV15 using RLT buffer (Qiagen RNeasy mini kit, 74104) supplemented with β-mercaptoethanol following manufacturer’s instructions. RNA was retrotranscribed using High-Capacity cDNA Reverse Transcription Kit (Thermofisher, 4368814). To determine the fold-enrichment of *Bcl6* cDNA in KD vs control conditions, LightCycler® 480 SYBR Green I Master kit (Roche) and comparative C_T_ method were used. The mRNA levels were normalized to *Rpl10* mRNA (housekeeping gene) (Table S3). Statistical analysis was performed using GraphPad Prism 9.

### Morphology analysis

#### Spine analysis

Neurolucida 360 and Neurolucida View Explorer were used to detect dendritic spines and perform spine analysis. Within Neurolucida 360, the backbone of a dendritic segment of 20-30 μm was tracked. 3-4 dendritic segments were reconstructed per animal. Further, spines were classified using Neurolucida and manually confirmed in the 3D environment based on the fluorescent signal. Neurolucida View Explorer was used to obtain quantitative spine measurements. Spine type distribution and spine density were assessed for control and KO/KD conditions. Statistical analysis was done using GraphPad Prism 10.

#### Mossy fiber bouton reconstructions

To reconstruct individual MFBs, aligned stacks were uploaded in Imaris (Bitplane). The number of filopodia was manually measured in 3D and only filopodia with a minimum length of 1.5 μm were counted. The 3D reconstructions of the MFB core were obtained by initially segmenting the volume containing the entire MF terminal. The segmented volume was subsequently masked and the MFB was reconstructed using the “New Surface” tool. Total surface area and volume of the MFB, and length of filopodia were determined for both control and KO/KD conditions through the ’Statistics’ function. 6-8 MFBs were reconstructed per animal. Statistical analysis was done using GraphPad Prism 10.

### Whole cell electrophysiology

For whole-cell patch clamp recordings, acute slices were prepared from P27-31 Rbp4-Cre:Rosa26-LSL-Cas9 animals injected with either lacZ gRNA (Control) or *Smad3* gRNAs (KO). In short, animals were anesthetized using isoflurane and after decapitation, the brain was quickly removed and transferred into ice-cold cutting solution (in mM): choline chloride 110, NaHCO3 26, Na-ascorbate 11.6, D-glucose 10, MgCl2 7, Na-pyruvate 3.1 , KCl 2.5, NaH2PO4 1.25, CaCl2 0.5; 300–315 mOsm, pH adjusted to 7.4, with 5% CO2/95% O2. Parasagittal slices (250 µm) containing hippocampus were made using a vibratome (VT1200, Leica Biosystems). Immediately after cutting the slices were transferred to 32-33 °C cutting solution for 6 min to recover and subsequently stored in holding solution (in mM): NaCl 126, NaHCO3 26, D-glucose 10, MgSO4 6, KCL 3, CaCl2 1, NaH2PO4 1, 300-310 mM, pH adjusted to 7.4, with 5% CO2/95% O2 at room temperature. Slices were stored for ∼1 hour before experiments. During experiments brain slices were continuously perfused in a submerged chamber (Warner Instruments) at a rate of 3-4 ml/min with recording solution (in mM) 127 NaCl, 2.5 KCl, 1.25 NaH2PO4, 25 NaHCO3, 1 MgCl2, 2 CaCl2, 25 D-glucose (pH 7.4 with 5% CO2/ 95% O2), including AP-5 (100 µM) and TTX (1 µM). All recordings were done between 31-33 °C. We used borosilicate glass recording pipettes (resistance 3.5–5 MΩ, Sutter P-1000) filled with internal solution (in mM): 126 CsMSF, 10 HEPES, 2.5 MgCl2, 4 ATP, 0.4 GTP, 10 Creatine Phosphate, 0.6 EGTA, 5 QX-314 and 3 mg/ml biocytin. Whole-cell patch-clamp recordings of granule cells in the dentate gyrus were done using either double EPC-10 amplifier under control of Patchmaster v2 x 32 (HEKA Elektronik) or Multiclamp 700B amplifier under control of Clampex 10.7 (Axon Instruments). Currents were recorded at 20 kHz and low-pass filtered at 3 kHz when stored. Series resistance was compensated to 70-80% and monitored throughout the experiments. For each granule cell both miniature excitatory synaptic inputs (mEPSCs; Vm= -90 mV) and miniature inhibitory synaptic inputs (mISPCs; Vm= 0 mV) were recorded. Miniature input was quantified using the Mini Analysis program (Synaptosoft).

### Read mapping

#### Single-nucleus Multiome read mapping

The generated raw fastq files were mapped to the *Mus musculus* (mm10) reference genome using the Cellranger Arc (v2.02) count function with intronic reads included by default.

#### RiboTag read mapping

The generated raw fastq files were mapped to the *Mus musculus* (mm10) reference genome using STAR v2.7.5b (*92*). Gene counts were then obtained using featureCounts function from SAMtools v1.10 (*93*).

#### Smart-seq2 read mapping

Adapter trimming was performed using the fastp tool (*94*) (Table S6). Reads were mapped to a custom Mus musculus mm10 reference genome, which included the tdTomato gene, using the STAR tool (*92*) (Table S6). Quality control was applied to remove low-quality cells, filtering out cells with low mapping percentages or those lacking tdTomato gene detection (Table S6).

### Sequencing data quality control

#### SS2 analysis

Data from different samples were merged and subjected to quality control and filtering. Specifically, cells with fewer than 500 genes, more than 8,000 genes, or more than 5% mitochondrial gene content were filtered out. Additionally, genes expressed in fewer than 6 cells were removed, and all mitochondrial genes, along with a list of sex-related genes (as referenced in (*47*), Table S7), were excluded from the resulting count matrix. This resulted in a total of 509 cells, with a median of 4276 genes detected per cell. The merged data was then normalized and log-transformed using the normalize_per_cell and log1p functions in Scanpy (v1.9.1) (*95*), respectively. Highly variable genes were identified using the ’seurat_v3’ method with n_top_genes = 4000, and the data was subsequently scaled. Finally, principal component analysis (PCA) was performed, and the first two principal components (PC1 and PC2) were visualized.

#### RiboTag analysis

First, for quality control, genes with extreme or low expression values were filtered out. Specifically, we required each gene to have an expression count greater than 50 in at least 3 samples, and no gene could have an expression value exceeding 500,000. Genes that did not meet these criteria were excluded from further analysis. Additionally, mitochondrial genes, sex-related genes (as referenced in (*47*), and genes associated with the GO term ’central nervous system myelination’ (GO:0022010)(*96*) were removed (Table S7).

Next, to estimate factors of unwanted variation, the RUVs function from the RUVSeq package v1.32.0 (*97*) was applied with k=2. Normalization for library size was performed, and the data was log2-transformed using the rlogTransformation function from the DESeq2 v1.38.3 package (*98*).

#### Multiome snRNA-seq analysis

Gene expression data quality control was performed independently for each sample using the Scanpy tool v1.9.1 (*95*). For each sample, genes expressed in at least three cells were retained, as well as cells expressing a minimum of 200 genes and with mitochondrial DNA content below 5%. Doublets were then identified and removed using the Scrublet tool (*99*), either by directly removing identified doublets or by excluding clusters with a high mean doublet score.

#### Merge Samples GEX multiome

Prior to merging the high-quality cells from the four different samples, mitochondrial genes and a list of sex-related genes (Table S7, (*47*)) were removed. Next, normalization and variance stabilization (*100*) were applied to each sample independently using the SCTransform() function from Seurat v4.3.0 (*101*) with the "glmGamPoi" method, regressing out mitochondrial gene percentage. Samples were then merged, and batch effects were corrected using 30 dimensions with Seurat and Harmony tool (*102*).

A total of 79,914 nuclei from four different time points were clustered using Louvain clustering (*103*) with a resolution of 0.9, resulting in 40 clusters. These clusters were manually annotated to cell types based on marker gene expression (Table S8).

For the focus of our analysis, cells belonging to the granule cell (GC) lineage—including neural progenitor cells (NPCs), type 1 and 2 neuroblasts, immature GCs (iGCs), and mature GCs (mGCs)—were identified and extracted based on GC lineage marker gene expression resulting in a total of 22,912 high-quality snRNA-seq cells. These cells were further filtered based on snATAC-seq quality (see next section for further details), leading to 18,907 cells, with a median of 3,499 unique molecular identifiers (UMIs) and 1,796 expressed genes per nucleus.

The selected cells (GC lineage) were then normalized using SCTransform() function from Seurat v4.3.0 (*101*) (applied separately by age). Samples were then combined and PCA was performed using all features. Next, non-linear dimensionality reduction was applied and then visualized using RunUMAP function from Seurat with the first 25 principal components used (number determined by inspecting elbow plot).

#### Gene expression comparison snMO versus SS2

To focus on transcriptionally active genes, for each timepoint in the SS2 data, we selected the top 40% of the most highly expressed genes based on their mean expression. The average expression profile of these selected genes was calculated for each cell type at the corresponding timepoint in the snMO snRNA-seq data. To quantify the similarity between gene expression profiles of SS2 and snMO, cosine similarity was computed using the cosine function from the SciPy package (version 1.12.0) (*104*).

#### Multiome snATAC-seq analysis

Each snATAC-seq sample was processed separately using the pycisTopic tool v2.0 (python implementation of cisTopic (*105*). First, using cell type annotations for each cell from snRNA-seq analysis, we generated pseudobulk profiles per cell type in GC lineage and then per age only for mGCs. Next, peaks were called on each pseudobulk profile using MACS2 v2.2.9.1 (*106*) with the parameters: --format BEDPE, --gsize mm, --qvalue 0.05, --keep-dup all, --shift 73, and --ext_size 146. To derive consensus peaks, we applied an iterative peak filtering approach (*107*) as implemented in pycisTopic v2.0, using a peak_half_width of 250 base pairs and the mm10 blacklist v2. This resulted in a total of 387,898 called peaks.

Quality control on consensus peaks was performed using Ensembl version 98 (https://sep2019.archive.ensembl.org/index.html) as the BioMart host for gene annotation. The pycisTopic qc function was applied separately to each sample. High-quality cells were retained based on the following criteria: a minimum TSS enrichment score of 5, FRIP score higher than 0.28, and a minimum number of 1,000 fragments. The different quality control criteria resulted in a total of 18,907 cells with high-quality snRNA-seq and snATAC-seq profiles. The 18,907 snATAC-seq nuclei yielded 257,570 accessible regions, with a mean of 4,118 accessible regions and 8,152 unique fragments per cell, a 53.13% fraction of reads in peaks (FRIP), and a median transcription start site (TSS) enrichment of 15.75.

For topic modeling, Latent Dirichlet Allocation (LDA) (*108*) using Gibbs sampling was performed, as implemented in pycisTopic, using Mallet v202108 with 500 iterations across models of [30, 40, 45, 50, 60, 65, 70, 75, 80, 85, 90, 95, 100] topics. Based on pycisTopic model selection metrics, we selected the model with 80 topics.

Next, we binarized the topic-region distributions using both Otsu thresholding (*109*) and the ntop method, selecting the top 3,000 regions. To impute dropouts, region-topic and topic-cell probabilities were multiplied with a scaling factor of 10^6, followed by normalization using a scaling factor of 10^4.

To identify differentially accessible regions (DARs), highly variable regions were extracted using the default parameters in pycisTopic. Cell-type-specific DARs were identified using the Wilcoxon test on the imputed accessibility matrix, with an adjusted p-value threshold of 0.01 and an absolute log fold change threshold of 1.

For comparisons across development, we identified DARs across different age groups of mGCs. This was done by comparing each age-specific group of mGCs against the remaining mGCs. Additionally, we performed pairwise Wilcoxon tests to identify DARs between each pair of ages in mGCs.

For DARs heatmap visualization, Signac package v1.9.0 (*110*) in R v4.2.2 was used. Chromatin accessibility data was first transformed by applying the Term Frequency-Inverse Document Frequency (TF-IDF) method (*111*) using the RunTFIDF() function on filtered high-quality cells. The average accessibility in mGCs of each age group was then plotted using ggplot2 v3.4.1 (*112*). Genes were linked to up- and down-regulated differentially accessible regions with a simple approach by identifying the closest gene to each region using the ClosestFeature() function from the Signac package (*110*). Gene Ontology (GO) enrichment analysis was then performed on the resulting gene sets using the enrichGO() function from the clusterProfiler package (*113*). The analysis used biological processes ontology, with p-value and q-value cutoffs of 0.05 and the Benjamini-Hochberg correction method. The top four enriched terms for each gene set were merged and visualized with a dot plot.

For dimensionality reduction of the snATAC-seq data using UMAP, Singular Value Decomposition (SVD) was performed using the RunSVD() function after identifying the top variable features (i.e., regions) with the FindTopFeatures() function, applying a minimum cutoff of 20. Finally, UMAP was computed using the first 30 dimensions derived from the Latent Semantic Indexing (LSI) reduction, with the first dimension discarded due to its correlation with sequencing depth. The snATAC-seq data was then visualized in UMAP two-dimensional space.

#### Pseudotime Inference SS2 GCs

Merged SS2 data was log-normalized using Seurat v4.3.0 (*101*) NormalizeData() function. The top 2000 variable features of the merged data were identified using the FindVariableFeatures() function, and the log-normalized data was scaled across all features using the ScaleData() function. To infer the pseudotime order of GCs, regularized ordinal regression models were fitted using ordinalNetCV() function from ordinalNet package v2.12 (*114*) with the ordered age as the response, using a cumulative link model (family = ’cumulative’). The model was evaluated with nFolds set to 5 and nFoldsCV set to 10. The regularization parameter alpha was optimized using 5-fold cross-validation, with the objective of minimizing the Mean Squared Error. To determine genes most associated with pseudotime, non-zero coefficients were extracted excluding any intercept terms. The prediction scores are generated by applying a linear combination of these selected genes expression values, with each gene weighted according to its corresponding coefficient.

### Pairwise differentially expressed genes: SS2 and RiboTag

**SS2** The normalized merged SS2 data was used with FindMarkers() function to perform pairwise differential gene expression using MAST test to compare each age group with the following age group. Genes were considered upregulated or downregulated if the adjusted p-value was less than 0.05 and the absolute log fold change exceeded 0.5. The numbers of up- and down-regulated genes between each pair of consecutive time points were visualized with a bar plot.

A generalized additive model (GAM) was fitted for each of the top 50 upregulated and downregulated genes of each pairwise comparison (ranked by log fold change) using the gam() function from the mgcv package v1.8.41 (*115*), with inferred pseudotime as the predictor and the family set to Gaussian. In order to define gene clusters, the Euclidean distance matrix of the GAM-fitted values was computed, and hierarchical clustering was performed using hclust() function from stats (v4.2.2, R) with Ward’s minimum variance method ("ward.D2") as the linkage criterion, with a cutoff height of 28.

GO enrichment analysis for biological processes (BP) was performed using the clusterProfiler v4.6.2 (*113*) package and the Mus musculus organism database (org.Mm.eg.db), with p-value and q-value cutoffs set to 0.05. For visualization, the top 3 GO terms from each analysis were extracted, merged, and visualized in a dot plot. Gene expression patterns of the top 50 upregulated and downregulated genes (Table S9) from each pairwise comparison were visualized in a heatmap (*116*) using the GAM-fitted expression values labeling only the top 20 genes per pairwise comparison that overlapped with genes from the SynGO database.

#### RiboTag

Pairwise differential expression analysis (comparing each age to the following age) was computed using the DESeq function from DESeq2, based on the negative binomial distribution and corrected for the estimated factors of unwanted variation (*97*, *98*). Shrinkage of effect sizes was applied using the lfcShrink function (with the "apeglm" method). Genes were considered differentially expressed if the adjusted p-value was less than 0.05 and the absolute log2 fold change exceeded 0.5. The numbers of up- and down-regulated genes between each pair of consecutive time points were visualized with a bar plot.

For clustering of differentially expressed genes (DEGs), the top 50 upregulated and downregulated DEGs per pairwise comparison (ranked by log2 fold change; Table S10) were selected and grouped based on their expression profiles using the degPatterns function from the DEGreport package (*117*) v1.39.6 with nClusters=6. Specifically, a distance matrix was created by calculating pairwise gene expression correlations using the mean expression per group, followed by clustering of genes using the DIANA method.

#### GRNs Inference

To identify gene regulatory networks (GRNs), the SCENIC+ algorithm was applied to the single nucleus multiome data using the scenicplus Snakemake pipeline from the scenicplus package v1.0a1 (*51*). Briefly, motif enrichment analysis was performed using pycisTarget v1.0a2 (*51*, *118*) on the region sets consisting of the identified DARs (see above) and binarized topic regions (from the Otsu and ntop methods), using the *Mus musculus* species and a custom ranking database using the consensus peaks identified for the GC lineage, with a padding of 1,000 base pairs. The motif similarity false discovery rate (FDR) threshold was set to 0.001, with an orthologous identity threshold of 0 and motif-transcription factor (TF) annotations based on the v10 clustering version (*51*).

pycisTarget analysis resulted in a set of cistromes modules consisting of TFs with their bound regions (regions that have significant enrichment of the TF motif). Regions in the resulting cistromes were then filtered to retain only those that were accessible in any mGC cluster. Gene annotations were obtained using the archived BioMart host v98 (September 2019). Finally, TF-to-gene and region-to-gene importance scores were inferred using a Gradient Boosting Machine (GBM) and with a search space defined as the larger of either the nearest gene boundary or 150 kb, and at least 1 kb upstream of the transcription start site (TSS) or downstream of the gene’s end.

Finally, enhancer-driven Regulons, in which TFs are linked to their target regions that are subsequently linked to their target genes, were constructed using a gene set enrichment analysis approach and then cell enrichment score of both eRegulon target gene and eRegulon target regions were calculated using AUCell algorithm (*118*). SCENIC+ inferred eRegulons were filtered to retain only activators (+/+, positive correlation of both TF expression-target region accessibility, and target region accessibility-target gene expression) and repressors (-/+, negative correlation of TF expression-target region accessibility and positive correlation of target region accessibility-target gene expression). For simplicity in this text, activators are referred to as TF(+) and repressors as TF(-) (rather than TF_+/+ and TF_-/+, respectively). In total 66 eRegulons were obtained.

PCA was performed on normalized and scaled eRegulons enrichment scores per cell (AUC eRegulons matrix using AUCell algorithm) using Scanpy package (*95*). We observed that the first principal components captured mainly the variability between cell types in the GC lineage rather than the variability between age groups (data not shown). Non-linear dimensionality reduction was then applied using UMAP with the first 8 principal components. We observed a separation of cells by their cell type following GC lineage across development from progenitors to the mature state.

#### Pseudotime Inference snMO mGCs

To infer pseudotime values for individual cells in snMO mGCs, we utilized eRegulon AUC scores, representing the enrichment of eRegulons per cell. The computed AUC scores were normalized per cell and log-transformed using a pseudocount of 1. Subsequently, Z-score normalization was applied to each eRegulon to center the activity scores to a mean of 0 and a standard deviation of 1 across all cells. The standardized values were then used to perform PCA via the prcomp() function from the stats package (v4.2.2, R). An elbow plot was generated to visualize the proportion of variance explained by each principal component in order to determine the optimal number of principal components (PCs) to retain for diffusion map analysis. A diffusion map of mGCs was created using the DiffusionMap() function from the destiny package v3.12.0 (*119*) with 12 PCs. mGCs were then clustered using the kmeans() function (stats package, v4.2.2, R) with 10 centers to obtain the trajectory starting cluster. mGCS maturation trajectory was subsequently inferred using the slingshot() function from the slingshot package v2.6.0 (*120*), with the diffusion map as the reduced dimensional space.

#### GC Lineage eRegulon Selection

To select cell type specific eRegulons across the GC lineage, two parallel selection methods were followed. eRegulons were first selected if their Spearman correlation value between eRegulon activity score and TF expression is higher than the mean correlation of all eRegulons. Next, eRegulons were also selected if they showed differential activity between two cell types (Wilcoxon rank sum test, adjusted p-value smaller than 0.0025 (Bonferroni corrected alpha), log fold change higher than 0.2 and active in at least 25% of the cells in that cell type). In addition, the eRegulon leading TF should also be differentially expressed in the same cell type (Wilcoxon rank sum test, adjusted p-value smaller than 0.0025 (Bonferroni adjusted), log fold change higher than 0.5 and active in at least 25% of the cells in that cell type). In total, 48 eRegulons were selected.

#### mGCs eRegulon Selection

To identify key eRegulons, two parallel approaches were used, focusing exclusively on cells annotated as mGCs. First, Spearman correlation was calculated between eRegulon activity scores (AUC values) and TF expression levels. eRegulons were selected if their correlation value exceeded the mean correlation across all eRegulons. Second, eRegulons were also chosen if they exhibited differential activity between two age groups, based on a Wilcoxon rank-sum test (adjusted p-value < 0.0033 (Bonferroni correction), log fold change > 0.2, and active in at least 0.25% of cells in that age group). Additionally, the corresponding TF had to be differentially expressed in the same age group (adjusted p-value < 0.0033 (Bonferroni correction), log fold change > 0.5, and expressed in at least 0.25% of cells). This analysis resulted in a total of 36 eRegulons active in mGCs. To identify and visualize different eRegulons clusters in mGCs, a linkage matrix of eRegulons was obtained by performing hierarchical clustering using the clustermap function from seaborn package v0.13.2 (*121*) with correlation as the metric for distance calculation. The dendrogram was then created using hierarchy cluster module from scipy v1.12.0 (*104*), with a color threshold (cut-off distance) set at 0.85.

PCA was performed on eRegulons AUC scores (enrichment per cell) in mGCs. This analysis revealed a clear separation of cells by age group, with cells following a continuous trajectory from the early time point (P5) to the late time point (P28).

GO enrichment analysis for eRegulon target genes was performed for biological process (BP) or cellular component (CC) using the clusterProfiler v4.6.2 (*113*) package and the *Mus musculus* organism database (org.Mm.eg.db), with p-value and q-value cutoffs set to 0.05. For visualization, the top 7 terms were selected, merged, and displayed in a dot plot.

eRegulon target region accessibility was visualized using Signac (v1.9.0) (*110*) in R v4.2.2. Region accessibility was first transformed by applying the Term Frequency-Inverse Document Frequency (TF-IDF) method (*111*) using the RunTFIDF() function on filtered high-quality cells. The average accessibility in mGCs for each age group was then plotted using ComplexHeatmap library v.2.14.0 (*116*). Regions were then labeled by their predicted target regions by SCENIC+. Signal plot of observed chromatin accessibility over specific target regions was shown using Seurat’s CoveragePlot() function with 1kbp (*Nptx1*), 500bp (*Tanc1*) and 5kbp (*Nptx1*), 75kbp (*Tanc1*) extension upstream, and downstream, respectively. SCENIC+ predicted target region-target gene links are also shown.

#### Correlation RiboTag GEX vs Multiome GEX

To compare gene expression across postnatal developmental timepoints in both the multiome and RiboTag data, the expression patterns of TFs and their predicted target genes were visualized for a subset of selected eRegulons. Specifically, the mean gene expression of eRegulons’ leading TF and target genes (normalized and log-transformed) per age group was calculated for both RiboTag RNA-seq and multiome snRNA-seq. These values were then Z-score normalized and visualized. Additionally, the ± standard deviation of target gene expression in RiboTag was calculated and visualized.

To assess the linear correlation in TFs gene expression between RiboTag RNA-seq and multiome snRNA-seq, the Pearson correlation coefficient was calculated. Specifically, the mean gene expression per age was used for this calculation, and the Pearson correlation was computed using the stats module from the SciPy v1.12.0 package (*104*). P-values were determined using Monte Carlo resampling with a uniform distribution for random sampling. However, given the relatively small sample size, these p-values may be subject to increased variability and potential inaccuracies and should be interpreted with caution. To quantify the linear correlation in eRegulon activity between RiboTag RNA-seq and multiome snRNA-seq, the Pearson correlation metric was also applied. Specifically, the Pearson correlation for each target gene was computed using the corrwith() function from pandasv1.5.0), and the mean correlation of all target genes was used to represent each eRegulon. The 95% confidence intervals for the Pearson correlation coefficients were estimated using the bootstrap() function from the scipy.stats module v1.12.0. A total of 1000 random paired resamples with replacement were generated to compute the confidence intervals.

## Acknowledgments

We thank Artemis Koumoundourou, Esther Klingler and Pierre Vanderhaeghen for critical reading of the manuscript, and De Wit lab and Aerts lab members for helpful discussion and comments. We are grateful to Gencay Kaan Polat for help with pilot experiments. We thank Lynette Lim (VIB – KU Leuven) for advice during single cell pilot experiments. We thank Aya Takeoka (VIB-Imec-KU Leuven Neuro Electronics Research Flanders) for sharing Rbp4-Cre and Ai9 mice. We thank Jan Cools (VIB – KU Leuven) for sharing Rosa26-LSL-Cas9 mice. We thank VIB Flow Core, VIB/CBD Bioimaging Core, VIB Nucleomics Core and VIB Single Cell Core for access to instruments and technical support.

## Funding

B.L-E. is supported by Fonds Wetenschappelijk Onderzoek (FWO, Belgium) PhD fellowship 1120821N and 1120823N, G.M. is supported by FWO PhD fellowship 11F1219N and 11F1221N; D.Das is supported by FWO Postdoctoral fellowship 12W5218N, 12W5221N and FWO Research Grant 1513320N; J.d.W. is supported by FWO Project Grants G0C4518N, G0A8720N and G0A8320N; FWO EOS Grant G0H2818N; ERANET-NEURON TAO2PATHY G0I3118N; FWO-SBO S005024N Enhancer-AI and a Methusalem Grant of KU Leuven/Flemish Government.

## Author contributions

B.L-E., D.Daa., and J.d.W. conceived the study and designed experiments. B.L-E. and D.Daa., and J.L. performed experiments and analyzed data. B.L-E., D.Daa., J.Vs., G.M., W.N., J.Vb., E.L., S.P., performed experiments. K.D. and S.A. provided guidance on bioinformatic analysis. S.P. and J.L. provided guidance on single cell/nuclei sample preparation. I.V., M.L., L.V. and K.W. performed electrophysiology experiments and analyzed data. B.L-E., D.Daa., and J.d.W. wrote the paper with input from all authors. All authors contributed to and approved the final version.

## Competing interests

J.d.W is scientific co-founder and served as scientific advisory board member of Augustine Tx. All other authors declare they have no competing interests.

## Data and materials availability

The data generated in this study will be available in the GEO database upon publication. Scripts used in this study will be available on Zenodo upon publication.

