## Supplementary figures and tables for "A dynamic gene regulatory code drives synaptic development of hippocampal granule cells"

**This PDF file includes:**

Figs. S1 to S8

Tables S1 to S5

Fig. S1

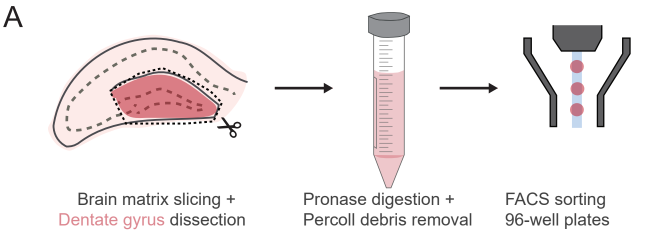


B

| Time points | #mice | sex | #cells |
|---|---|---|---|
| P5 | 1 | F | 94 |
| P7 | 1 | M | 76 |
| P10 | 1 | F | 91 |
| P15 | 1 | M | 93 |
| P28_E1 | 1 | F | 88 |
| P28_E2 | 1 | M | 67 |
| Total | | | 509 |

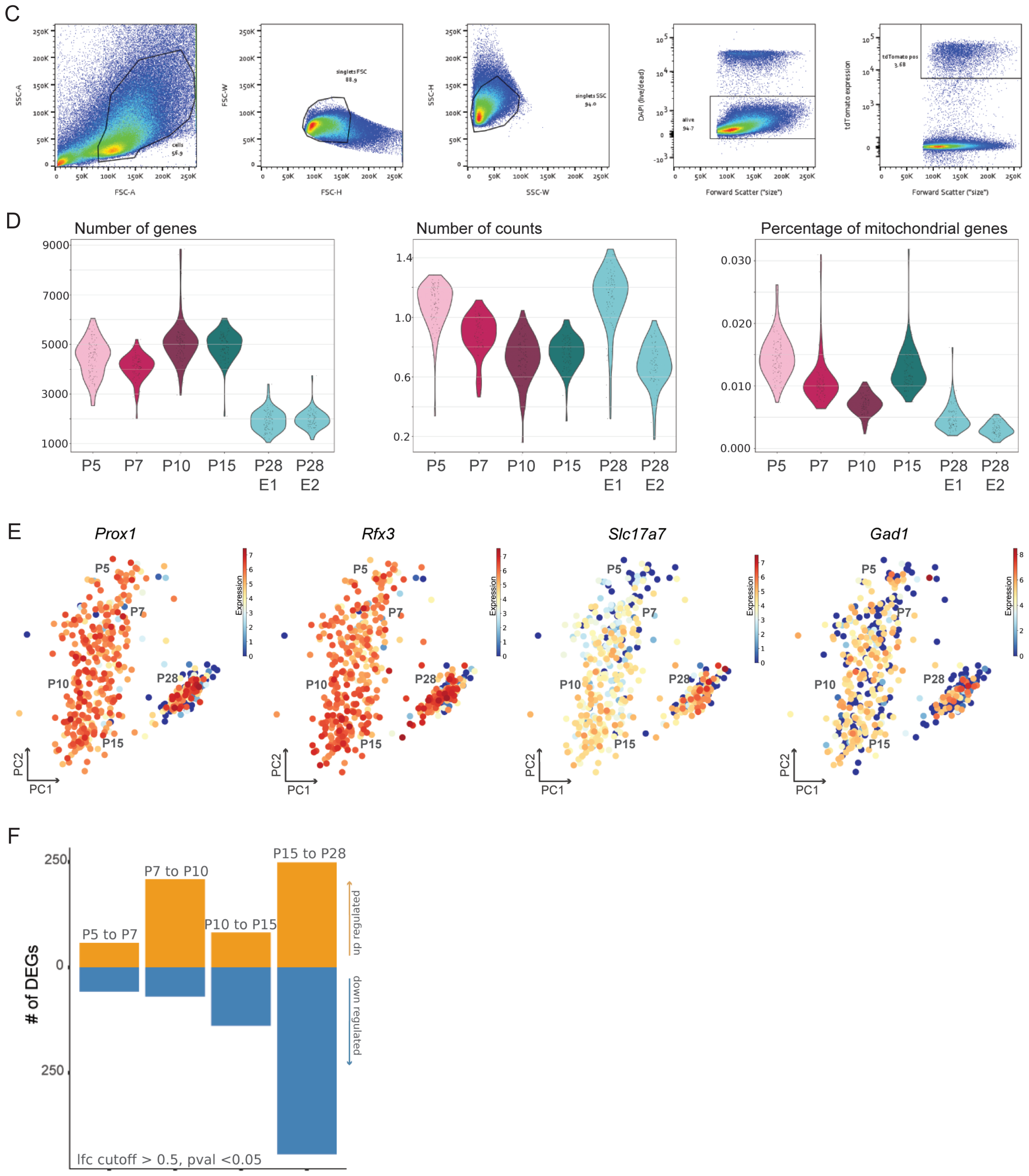

**Fig. S1. Methodology and quality control of single-cell RNA sequencing (scRNAseq) experiments. A)** Methodology for scRNA sequencing experiments. Brain slices from Rbp4-Cre:Ai9 mice were made using a 500 µm brain matrix, and the dentate gyrus (DG) was dissected. Next, the DG was chopped in smaller pieces and digested using Pronase. Following the digestion, the tissue pieces were homogenized using two-size Pasteur pipettes and the mix was loaded in a 20% Percoll solution to remove debris. Following centrifugation, the pellet was resuspended in OptiMEM-trehalose, and DAPI was added. Fluorescence Activated Cell Sorting (FACS) was done based on DAPI and tdTomato signal, into 96-well plates containing lysis buffer (see Methods for details). **B)** Table shows number of animals per timepoint, sex of the animal and number of cells obtained. One animal per timepoint except for P28 where two experiments (E) were performed. Three female (F) and three male (M) mice were used in total. On average 85 cells were obtained per experiment, with a total of 509 cells post-quality control. **C)** FACS gating strategy to select cells, singlets and tdTomato-positive and DAPI-negative signal. **D)** Quality control of the scRNAseq shows number of genes, number of counts and percentage of mitochondrial genes obtained per timepoint and experiment. **E)** PCA colored by the expression of known granule cell and neuronal marker genes over time, i.e. *Prox1*, *Rfx3*, *Slc17a7* and *Gad1*. **F)** Bar plot represents number of differentially expressed genes (DEGs) from pairwise comparisons (log-fold change (lfc) >0.5 and adjusted p-value <0.05). 58 genes up- and 57 down-regulated from P5 vs P7 comparison, 209 genes up- and 69 down-regulated from P7 vs P10 comparison, 83 genes up- and 138 genes down-regulated from P10 vs P15 comparison, and 249 genes up- and 443 genes down-regulated from P15 vs P28 comparison (results listed in table S9).

Fig. S2

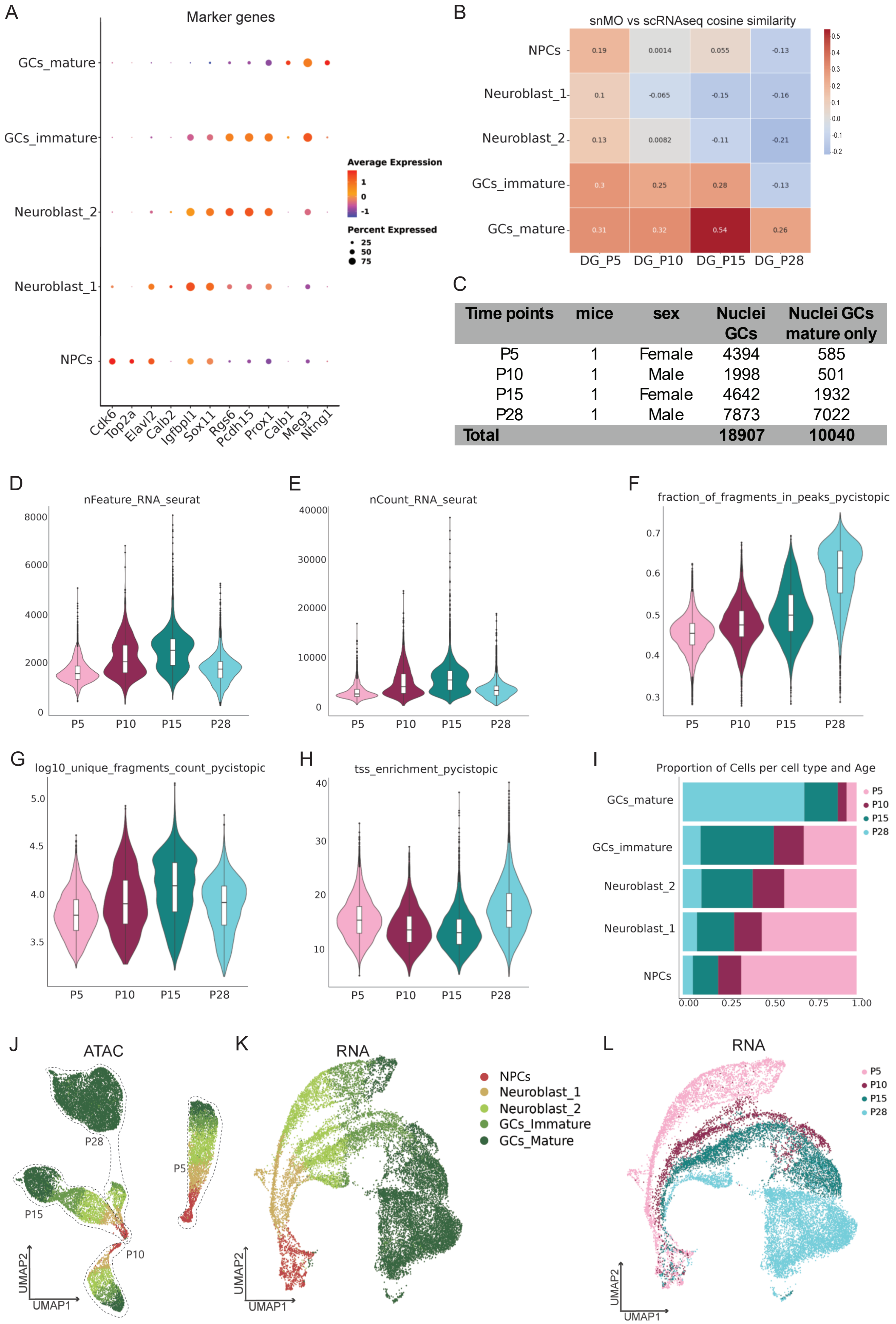


| Time points | mice | sex | Nuclei GCs | Nuclei GCs mature only |
|---|---|---|---|---|
| P5 | 1 | Female | 4394 | 585 |
| P10 | 1 | Male | 1998 | 501 |
| P15 | 1 | Female | 4642 | 1932 |
| P28 | 1 | Male | 7873 | 7022 |
| **Total** | | | **18907** | **10040** |

**Fig. S2. Quality control snMO experiments. A)** Dot plot shows marker genes across all cell types from the GC lineage (GC mature (mGCs), GC immature, Neuroblasts type 1 and 2, and Neuronal Progenitor Cells (NPCs)) (Table S8). **B)** Heatmap shows cosine similarity for Rbp4-Cre:Ai9 cells from scRNAseq dataset (labeled as DG_P5, DG_P10, DG_P15, DG_P28) vs DG cell types from snMO dataset (labelled as NPCs, Neuroblasts_1, Neuroblasts_2, GCs_immature, GCs_mature). **C)** Table shows number of animals per timepoint, sex of the animal and number of nuclei obtained from the snMO experiments. One animal per timepoint, two female and two male mice were used in total. A total of 18907 nuclei were obtained, with 10040 being mGCs. **(D-H)** Violin plots show **D)** number of genes per nuclei, **E)** number of counts per nuclei, **F)** fraction of fragments in peaks, **G)** unique fragments count, and **H)** transcription start site (tss) enrichment across all timepoints (P5, P10, P15, P28). **I)** Bar plot shows the proportion of nuclei per cell type across all timepoints. **J)** UMAP dimensionality reduction of GC lineage based on snMO chromatin accessibility profiles and colored by cell type. **(K, L)** UMAPs of GC lineage based on snMO gene expression profiles and colored by cell type **K)** and timepoint **L)**.

Fig. S3

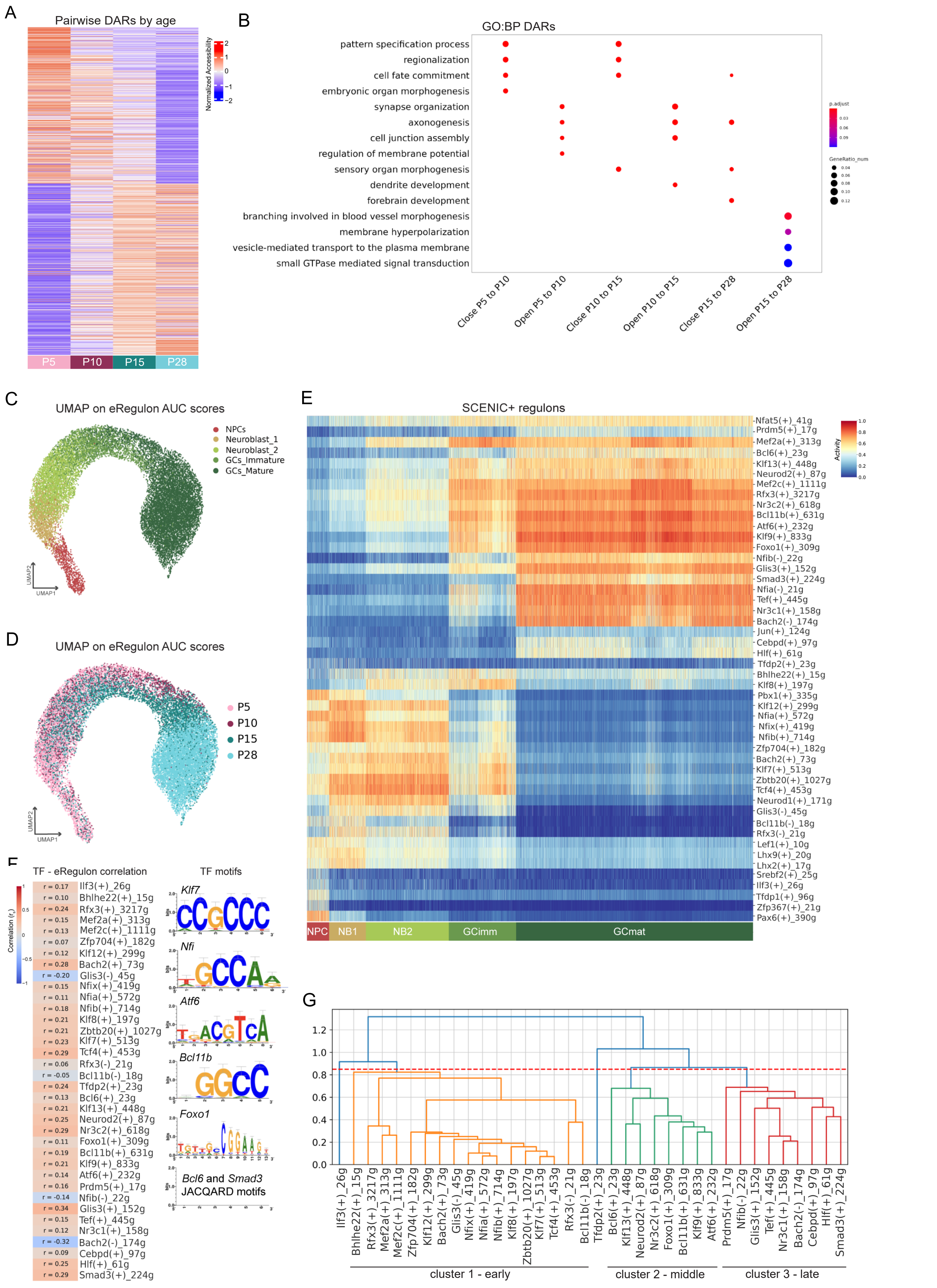
A
Pairwise DARs by age
Normalized Accessibility
P5
P10
P15
P28
B
GO:BP DARs
pattern specification process
regionalization
cell fate commitment
embryonic organ morphogenesis
synapse organization
axonogenesis
cell junction assembly
regulation of membrane potential
sensory organ morphogenesis
dendrite development
forebrain development
branching involved in blood vessel morphogenesis
membrane hyperpolarization
vesicle-mediated transport to the plasma membrane
small GTPase mediated signal transduction
Close P5 to P10
Open P5 to P10
Close P10 to P15
Open P10 to P15
Close P15 to P28
Open P15 to P28
p.adjust
GeneRatio_num
C
UMAP on eRegulon AUC scores
NPCs
Neuroblast_1
Neuroblast_2
GCs_Immature
GCs_Mature
D
UMAP on eRegulon AUC scores
P5
P10
P15
P28
E
SCENIC+ regulons
Activity
NPC
NB1
NB2
GCimm
GCmat
F
TF - eRegulon correlation
Correlation (r_s)
TF motifs
Klf7
Nfi
Atf6
Bcl11b
Foxo1
Bcl6 and Smad3
JACQARD motifs
G
cluster 1 - early
cluster 2 - middle
cluster 3 - late

**Fig. S3. Differentially accessible regions (DARs) in mature granule cells (mGCs) and SCENIC+ results. A)** Heatmap of mean normalized accessibility of differentially accessible regions (DARs) obtained by pairwise analysis across all timepoints. **B)** Dot plot represents gene ontology (GO) biological process (BP) terms of DAR-associated genes (closest gene per region). **(C, D)** UMAP dimensionality reduction of GC lineage nuclei based on eRegulon AUC (enrichment per cell) scores and labelled by **C)** cell type and **D)** timepoint. **E)** Heatmap of SCENIC+ predicted eRegulon AUC scores across the GC lineage: neural progenitor cells (NPCs), neuroblasts type 1 and 2, GC immature and GC mature. **F)** Heatmap of Spearman correlation ($r_s$) between leading TF expression-eRegulon activity in mGCs. The motifs of selected TFs are also shown. **G)** Dendrogram shows clustering analysis of eRegulons active in mGCs (Fig. 2D), cut off threshold shown as red line (0.85).

Fig. S4

A

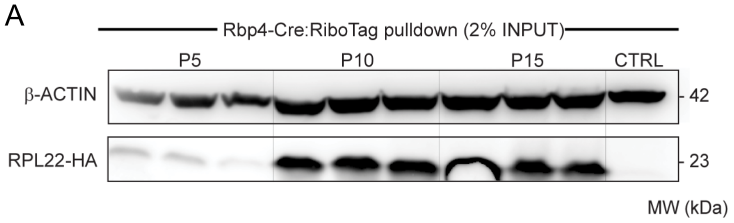


B

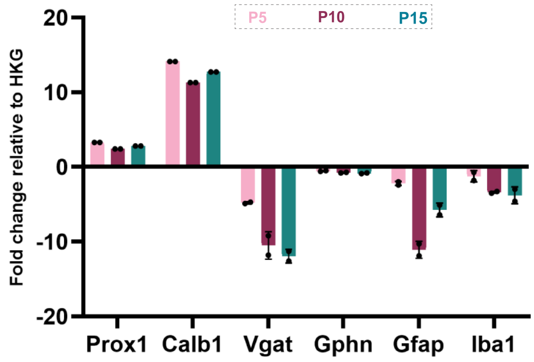


C

| Time points | #hippocampi | samples | sex |
|---|---|---|---|
| P5 | 3-4 | 5 | M, F |
| P7 | 4 | 3 | M, F |
| P10 | 2 | 3 | M, F |
| P15 | 2 | 3 | M, F |
| P21 | 2 | 3 | M, F |
| P28 | 2 | 3 | M, F |

D

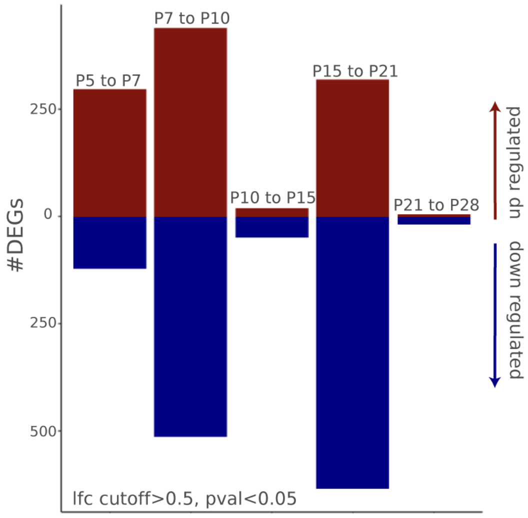


E

F

G

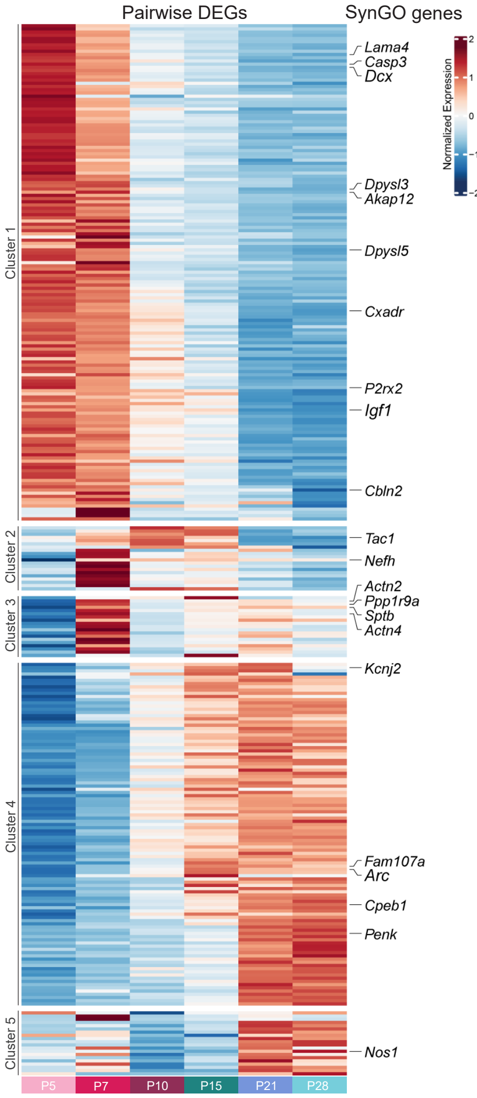


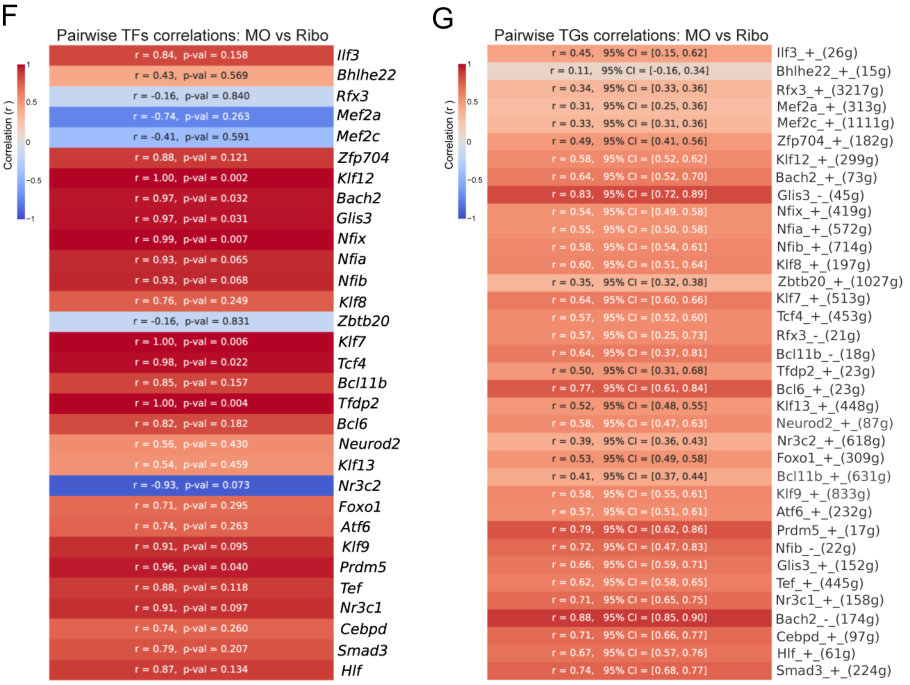

**Fig. S4. RiboTag dataset quality control and characterization. A)** Western blot shows RPL22-HA and β-ACTIN protein levels in 2% input Rbp4-Cre:Ribotag or control samples ($RiboTag^{Tg/Tg}$). **B)** RT-qPCR results show fold change relative to *Pkg1* housekeeping gene (HKG) for GC marker genes *Prox1*, *Calb1*; vesicular GABA transporter gene *Vgat*; inhibitory postsynaptic scaffold gene *Gphn*; astrocyte marker gene *Gfap* and microglia marker gene *Iba1*, at P5, P10 and P15. **C)** Number of hippocampi used per sample, and number of samples sequenced per timepoints. Animals from both sexes were used (F, female; M, male). **D)** Bar plot represents number of differentially expressed genes (DEGs) from pairwise comparisons across timepoints (log-fold change (lfc) >0.5 and adjusted p-val <0.05). 327 genes up- and 147 down-regulated from P5 vs P7 comparison, 442 genes up- and 464 down-regulated from P7 vs P10 comparison, 29 genes up- and 63 genes down-regulated from P10 vs P15 comparison, 396 genes up- and 719 genes down-regulated from P15 vs P21 comparison, and 6 genes up- and 17 genes down-regulated from P21 vs P28 comparison (results listed in table S10). **E)** Heatmap of the five clusters obtained from the top 50 DEGs of pairwise comparisons between timepoints. SynGO-annotated genes within the top 20 genes of each pairwise comparison are highlighted. **F)** Heatmap shows pairwise comparisons (Pearson correlation) for transcription factor (TF) expression in single-nuclei multiome (snMO) vs RiboTag datasets for TFs obtained in SCENIC+ analysis (Fig. 2D). **G)** Heatmap shows pairwise correlations (Pearson correlation) for target gene (TG) expression in snMO vs RiboTag for TGs of each eRegulon obtained in SCENIC+ (Fig. 2D).

Fig. S5

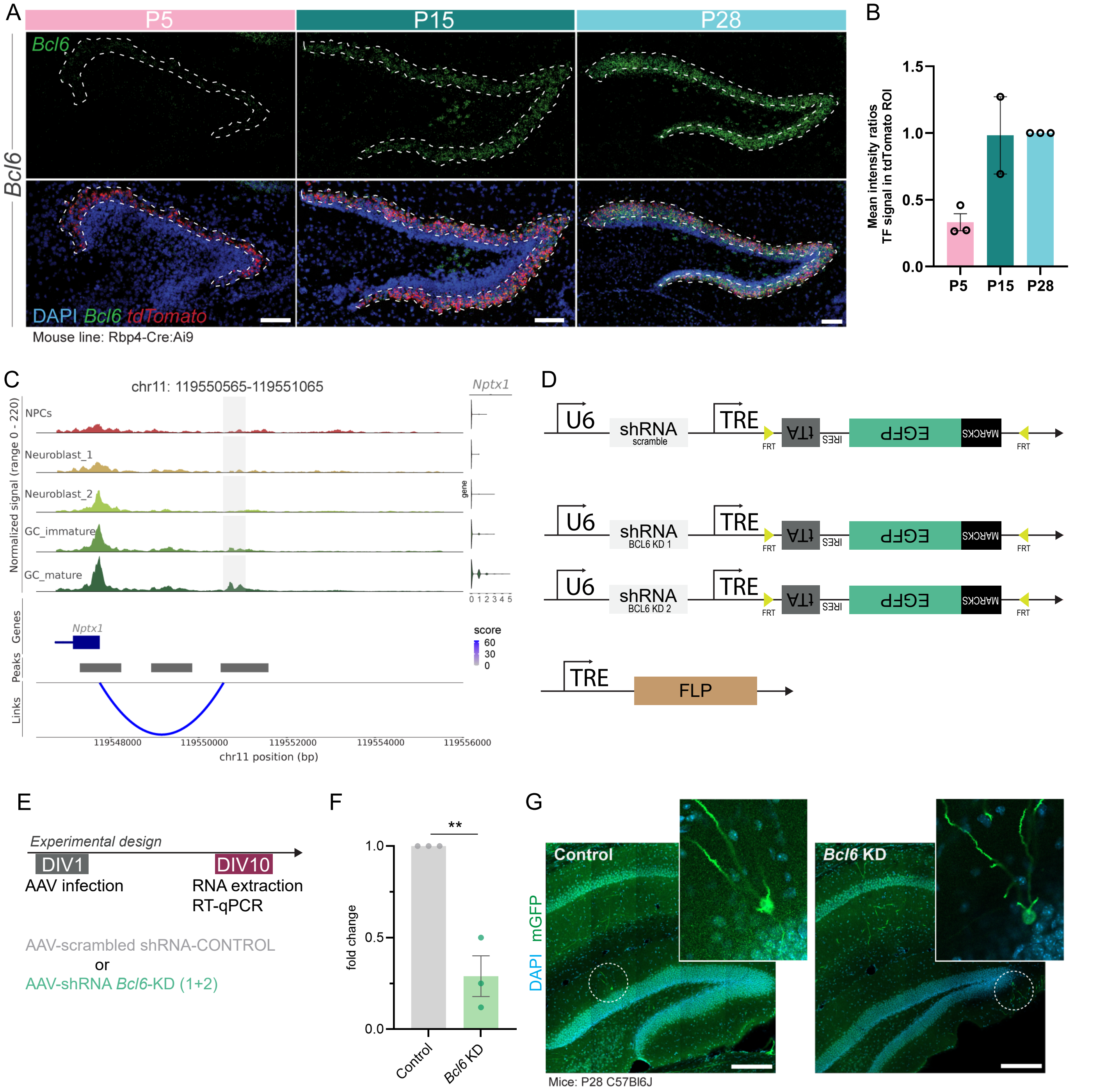

A
P5
P15
P28
Bcl6
DAPI Bcl6 tdTomato
Mouse line: Rbp4-Cre:Ai9
B
Mean intensity ratios TF signal in tdTomato ROI
1.5
1.0
0.5
0.0
P5
P15
P28
C
chr11: 119550565-119551065
Nptx1
Normalized signal (range 0 - 220)
NPCs
Neuroblast_1
Neuroblast_2
GC_immature
GC_mature
0 1 2 3 4 5
Genes
Peaks
Links
score
60
30
0
119548000
119550000
119552000
119554000
119556000
chr11 position (bp)
D
U6
shRNA
scramble
TRE
FRT
tTA
IRES
EGFP
MARCKS
shRNA
BCL6 KD 1
shRNA
BCL6 KD 2
FLP
E
Experimental design
DIV1
AAV infection
DIV10
RNA extraction
RT-qPCR
AAV-scrambled shRNA-CONTROL
or
AAV-shRNA Bcl6-KD (1+2)
F
**
fold change
1.0
0.5
0.0
Control
Bcl6 KD
G
Control
Bcl6 KD
DAPI mGFP
Mice: P28 C57Bl6J

**Fig. S5. *Bcl6* characterization, experimental design and in vitro validation. A)** RNAscope for *Bcl6* and *tdTomato* transcripts at P5, P15 and P28 in Rbp4-Cre:Ai9 animals. *Bcl6* transcripts (green) shown above, and their colocalization with tdTomato (red) and DAPI (blue) is shown below. Dotted outline marks region of *tdTomato*-expressing GCs. Scale bars 100 μm. **B)** RNAscope quantification of *Bcl6* signal in *tdTomato* region of interest (ROI) over time, normalized to P28 values. n=2-3 animals per timepoint. Bar represents mean±SEM, each dots represents an animal. **C)** Coverage plot for *Nptx1* region across timepoints for granule cell (GC) lineage (neural progenitor cells (NPCs), neuroblasts type 1 and 2, GC immature and GC mature). *Nptx1* region is located on chromosome (chr) 11 between base pairs (bp) 119550565 and 119551065 (grey shade). Arcs represent SCENIC+ predicted region-gene links. **D)** Schematic of plasmids pAAV-U6-TRE-fDIO-mGFP-IRES-tTA containing scrambled shRNA, *Bcl6* shRNA 1, or *Bcl6* shRNA 2, and pAAV-TRE-FLP used for Supernova sparse labeling. TRE: tetracyclin responsive element, IRES: internal ribosomal entry site, tTA: tetracycline transactivator, MARCKS: myristoylated alanine-rich protein kinase C substrate. FRT (Flp recognition target) sites are indicated with yellow arrowheads. **E)** Experimental design in vitro validation of *Bcl6* shRNAs. C57Bl6J (wild type) mouse primary cortical cultures were infected at day in vitro (DIV)1 with AAVs harboring control scrambled shRNA or a 1:1 mix of *Bcl6* shRNAs. RNA was extracted for RT-qPCR analysis at DIV10. **F)** RT-qPCR results for *Bcl6* transcript levels in *Bcl6* KD vs control cultures (fold-change relative to *Rpl0* housekeeping gene; n=3 independent cultures, t-test, p-value = 0.0031). Bar represents mean±SEM, each dots represents an independent culture. **G)** Overview images of the DG of animals injected with AAVs harboring control or *Bcl6* shRNAs. Insets: mGFP-labeled GCs in the outer part of the GC layer. mGFP: membrane-bound green fluorescent protein. Scale bars 200 μm.

Fig. S6

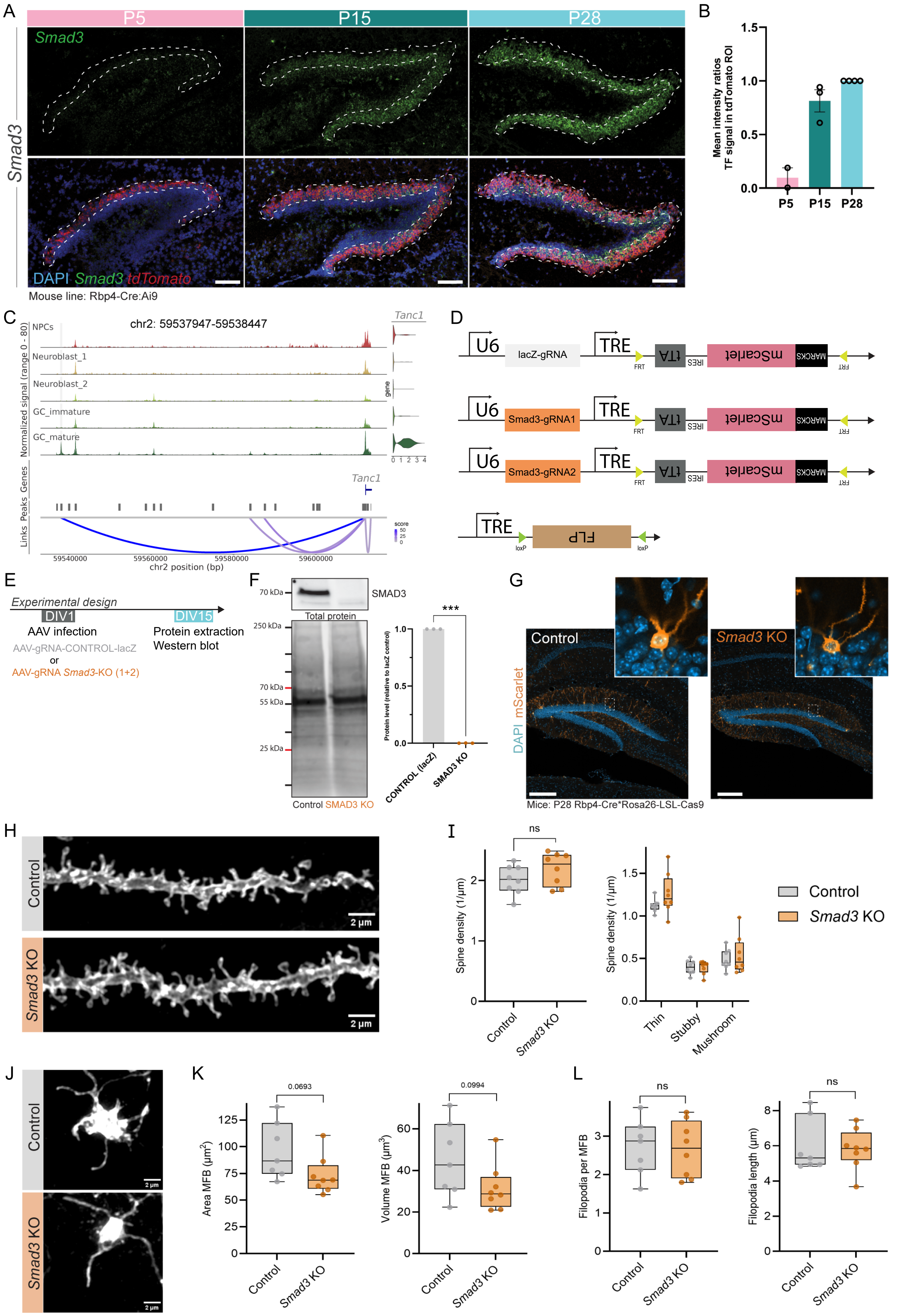
A
P5
P15
P28
Smad3
DAPI Smad3 tdTomato
Mouse line: Rbp4-Cre:Ai9
B
Mean intensity ratios TF signal in tdTomato ROI
1.5
1.0
0.5
0.0
P5
P15
P28
C
chr2: 59537947-59538447
NPCs
Neuroblast_1
Neuroblast_2
GC_immature
GC_mature
Normalized signal (range 0 - 80)
Tanc1
Genes
Peaks
Links
score
59540000
59560000
59580000
59600000
chr2 position (bp)
D
U6
lacZ-gRNA
TRE
FRT
tTA
IRES
mScarlet
MARCKS
Smad3-gRNA1
Smad3-gRNA2
FLP
loxP
E
Experimental design
DIV1
DIV15
AAV infection
Protein extraction
Western blot
AAV-gRNA-CONTROL-lacZ
or
AAV-gRNA Smad3-KO (1+2)
F
70 kDa
SMAD3
Total protein
250 kDa
70 kDa
55 kDa
25 kDa
Control SMAD3 KO
***
Protein level (relative to lacZ control)
1.0
0.5
0.0
CONTROL (lacZ)
SMAD3 KO
G
Control
Smad3 KO
DAPI mScarlet
Mice: P28 Rbp4-Cre*Rosa26-LSL-Cas9
H
Control
Smad3 KO
2 µm
I
ns
Spine density (1/µm)
2
1
0
Control
Smad3 KO
1.5
1.0
0.5
0.0
Thin
Stubby
Mushroom
Control
Smad3 KO
J
Control
Smad3 KO
2 µm
K
0.0693
Area MFB (µm²)
125
100
75
50
25
0
Control
Smad3 KO
0.0994
Volume MFB (µm³)
60
40
20
0
L
ns
Filopodia per MFB
3
2
1
0
ns
Filopodia length (µm)
8
6
4
2
0

**Fig. S6. Loss of *Smad3* in GCs does not alter dendritic spines or MFB morphology. A)** RNAscope for *Smad3* and *tdTomato* transcripts at P5, P15 and P28 in Rbp4-Cre:Ai9 animals. *Smad3* transcripts (green) shown above, and their colocalization with *tdTomato* (red) and DAPI (blue) is shown below. Dotted outline marks region of *tdTomato*-expressing GCs. Scale bars 100 μm. **B)** RNAscope quantification of *Smad3* signal in *tdTomato* region of interest (ROI) over time, normalized to P28 values. n=2-4 animals per timepoint. Bar represents mean±SEM, each dots represents an animal. **C)** Coverage plot for *Gabra1* region across timepoints for GC lineage (neural progenitor cells (NPCs), neuroblasts type 1 and 2, GC immature and GC mature). *Tanc1* region is located on chromosome (chr) 2 between base pairs (bp) 59537947 and 59538447 (grey shade). Arcs represent SCENIC+ predicted region-gene links. **D)** Schematic of plasmids pAAV-U6-TRE-fDIO-mScarlet-IRES-tTA containing *lacZ* gRNA or *Smad3* gRNAs, and pAAV-TRE-DIO-FLP. TRE: tetracyclin responsive element, IRES: internal ribosomal entry site, tTA: tetracycline transactivator, MARCKS: myristoylated alanine-rich protein kinase C substrate. FRT sites are indicated with yellow arrowheads; loxP sites are indicated with green arrowheads. **E)** Experimental design in vitro validation of *Smad3* gRNAs. H11-Cas9 mouse primary cortical cultures were infected at day in vitro (DIV)1 with control AAV or a 1:1 mix of AAVs harboring *Smad3* gRNAs. Protein was extracted for western blot analysis at DIV15. **F)** Left panels, representative western blot for SMAD3 and total protein stain on control and *Smad3* KO samples. Right graph, quantification of SMAD3 levels relative to total protein levels and normalized to control cultures (n=3 independent cultures, t-test, p-value < 0.0001). Bar represents mean±SEM, each dots represents an independent culture. **G)** Overview image of the DG of mice injected with AAVs harboring control or *Smad3* gRNAs. Insets: mScarlet-labeled GCs in the outer part of the GC layer. mScarlet: membrane-bound Scarlet. Scale bars 200 μm. **H)** Representative images of dendritic segments of GCs from the medial molecular layer of the DG for control (grey) and *Smad3* KO animals (orange). **I)** Spine analysis shows non-significant (ns) changes in spine density or morphology in *Smad3* KO animals compared to controls (n=4 20 μm dendrite segments analyzed per animal; n=8 mice for control; n=8 mice for *Smad3* KO; nested t-test). **J)** Representative images of MFBs from control (grey) or *Smad3* KO animals (orange). **K)** MFB analysis shows a non-significant decrease in area and volume of MFBs in *Smad3* KO animals compared to controls (6-8 MFBs analyzed per animal, n=7 mice for control; n=8 mice for *Smad3* KO; nested t-test). **L)** MFB structure analysis shows non-significant changes in filopodia number and length (Nested t-test). Box-and-whisker plots in I, K and L show median, interquartile range, minimum, and maximum; with each dot representing one animal.

Fig. S7

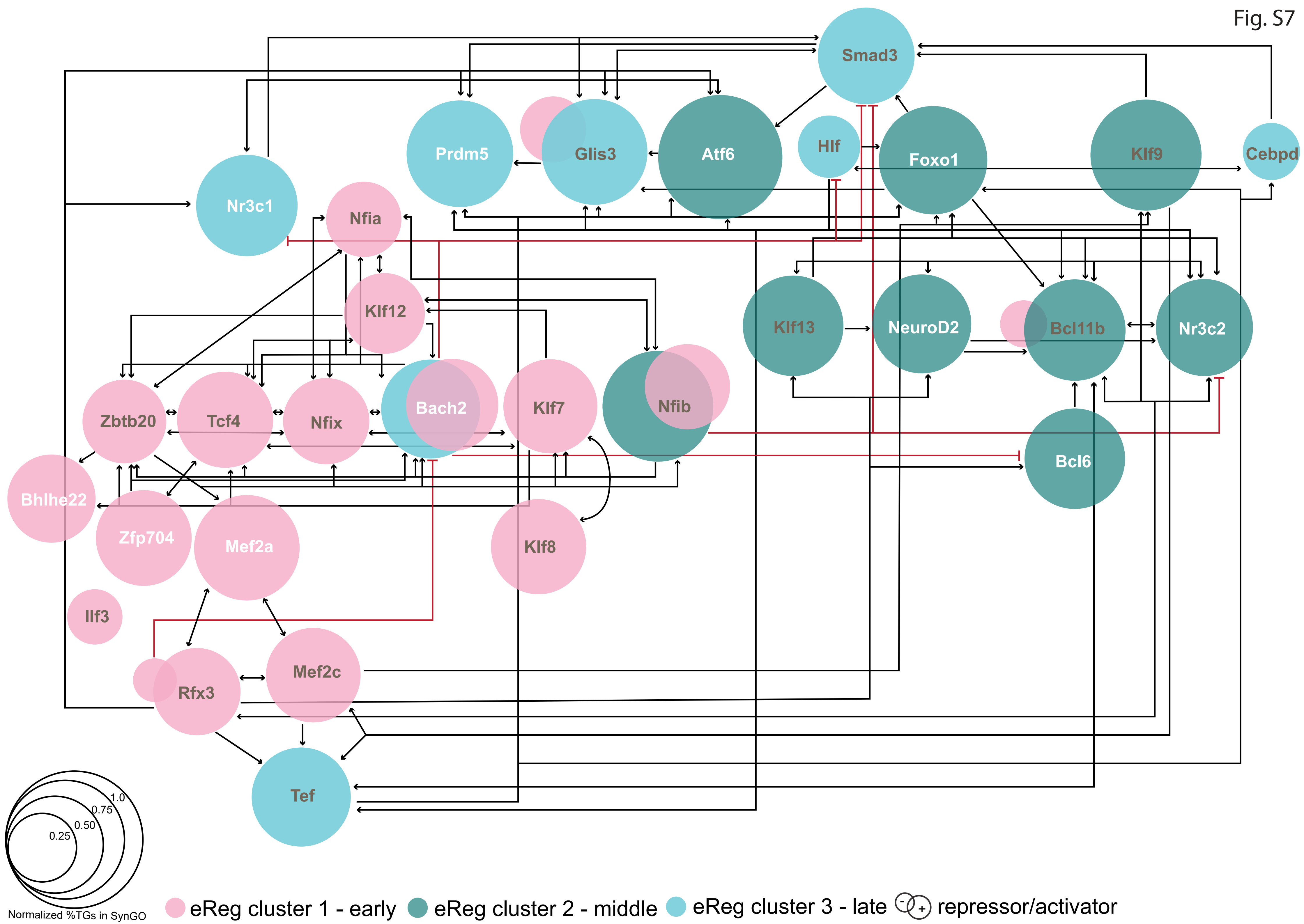

Smad3
Prdm5
Glis3
Atf6
Hlf
Foxo1
Klf9
Cebpd
Nr3c1
Nfia
Klf12
Klf13
NeuroD2
Bcl11b
Nr3c2
Zbtb20
Tcf4
Nfix
Bach2
Klf7
Nfib
Bcl6
Bhlhe22
Zfp704
Mef2a
Klf8
Ilf3
Rfx3
Mef2c
Tef
1.0
0.75
0.50
0.25
Normalized %TGs in SynGO
eReg cluster 1 - early
eReg cluster 2 - middle
eReg cluster 3 - late
repressor/activator

**Fig. S7. Schematic representation of GRN model.** GRN model of postnatal GC development showing eRegulons and the predicted interactions of leading TFs (incoming and outgoing; activating (black lines) and repressing (red lines)). eRegulons are represented by circles, colored depending on the cluster the eRegulon belongs to (cluster 1-early, pink; cluster 2-middle, green; cluster 3-late, blue; see also Fig. 2D, Fig. S3G). The size of the circles for each eRegulon is determined based on the percentage of SynGO-annotated synaptic target genes (TGs), normalized to the eRegulon with the highest proportion of SynGO-annotated synaptic genes, Atf6(+). eRegulons with leading TFs that act as both activators and repressors are represented by two circles, with the left circle representing the repressor, and the right circle representing the activator. TFs labeled in grey also target themselves (e.g. *Mef2c*), while those labeled in white do not (e.g. *Mef2a*).

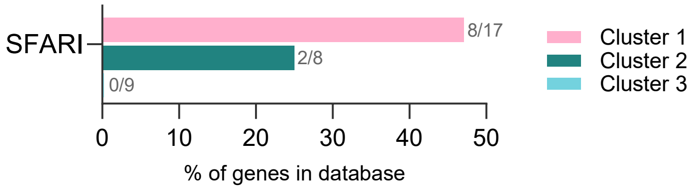


**Fig. S8. Percentage of leading transcription factors (TFs) detected in SFARI per cluster.** The percentage of cluster 1 genes is represented in pink, of cluster 2 in green, and of cluster 3 in blue.

Table S1

| **Leading TF** | **Known role in the DG** (pubmed search: TF + dentate gyrus) | **Known DG expression** https://developingmouse.brain-map.org/ (checking only P4-P14-P28) https://knowledge.brain-map.org/ (adult) | **Listed in Rasetto et al, 2024 Sci Advances** https://pubmed.ncbi.nlm.nih.gov/39028813/ | **Link to neurodevelopmental disorders (NDD)** https://genetrek.pasteur.fr/ https://gene.sfari.org/ |
|---|---|---|---|---|
| Ilf3 | No | Gene not found | No | No results |
| Bhlhe22 | No | Yes, from P4 | Regulon 3rd state<br>TF expressed neuroblasts + GC imm | Candidate NDD gene |
| Rfx3 | No | Yes, adult brain | No | Autism-associated genes SFARI |
| Mef2a | Adult hippocampal neurogenesis https://pubmed.ncbi.nlm.nih.gov/22542682/ https://pubmed.ncbi.nlm.nih.gov/26286136/ | Not expressed in DG | No | No results |
| Mef2c | Loss of Mef2c impairs learning and increases number of excitatory synapses https://pubmed.ncbi.nlm.nih.gov/18599438/ | Yes, from P14 | TF expressed in GCimm + GCmat | Autism-associated genes SFARI |
| Zfp704 | No | Yes, adult brain | No | No results |
| Klf12 | No | Yes, adult brain | Regulon 3rd state<br>TF expressed neuroblasts + GC imm | No results |
| Bach2 | No | Yes, adult brain | No | Candidate NDD genes |
| Glis3 | No | Yes, from P14 | TF in RGL and GCmat | High Confidence NDD gene |
| Nfix | Abnormal hippocampal morphology in het Nfix mice + memory deficits | Yes, from P4 | No | Autism-associated genes SFARI |

| | | | | |
|---|---|---|---|---|
| | https://pubmed.ncbi.nlm.nih.gov/23776487/<br>Smaller DG than controls<br>https://pubmed.ncbi.nlm.nih.gov/37283649/ | | | |
| Nfia | Smaller DG upon loss of Nfia<br>https://pubmed.ncbi.nlm.nih.gov/32166136/ | Yes, from P4 | No | Autism-associated genes<br>SFARI |
| Nfib | Smaller DG upon loss of Nfib<br>https://pubmed.ncbi.nlm.nih.gov/32166136/ | Yes, from P4 | No | Autism-associated genes<br>SFARI |
| Klf8 | No | Yes, adult brain | TF expressed in GC immature | No results |
| Zbtb20 | Loss of Zbtb20 affects DG morphology during early embryonic development<br>https://pubmed.ncbi.nlm.nih.gov/22689450/ | Yes, from P4 | No | Autism-associated genes<br>SFARI |
| Klf7 | No | Yes, from P4 | Regulon 3rd state | Autism-associated genes<br>SFARI |
| Tcf4 | Absence of Tcf4 in dentate neuroepithelium leads to abnormal progenitor migration<br>https://pubmed.ncbi.nlm.nih.gov/31845732/ | Yes, from P4 | No | Autism-associated genes<br>SFARI |
| Bcl11b | Regulates synaptic vesicle recruitment and long-term potentiation at MF synapses<br>https://pubmed.ncbi.nlm.nih.gov/38358390/ | Yes, from P4 | TF from GCimm to Gcmat | High Confidence NDD gene<br>Autism-associated genes<br>SFARI |
| Tfdp2 | No | Yes, from P14 | No | No results |
| Bcl6 | No | Yes, from P14 | Regulon 4th state<br>TF in RGL and GCmat | Candidate NDD gene |

| | | | | |
|---|---|---|---|---|
| Klf13 | No | Yes, adult | TF GC young and GCmat | No results |
| Neurod2 | Increased Neurod2 expression upon hypoxia https://pubmed.ncbi.nlm.nih.gov/26899848/ Promotes DG neurogenesis https://pubmed.ncbi.nlm.nih.gov/28740463/ Increased Neurod2 levels in the DG upon seizures https://pubmed.ncbi.nlm.nih.gov/11750069/ | Yes, from P4 | TF NBs, GC imm, GC young and GC mat | No results |
| Nr3c2 | Expression in DG not altered upon maternal separation https://pubmed.ncbi.nlm.nih.gov/33165192/ Antagonist of Nr3c2 reduces vGluT2 expression in the DG https://pubmed.ncbi.nlm.nih.gov/39237618/ | Yes, from P4 | No | Autism-associated genes SFARI |
| Foxo1 | Phosphorylation of Foxo1 increased in the DG upon exercise https://pubmed.ncbi.nlm.nih.gov/19936256/ Foxo1 DG expression downregulated in NF-kB deficient mice https://pubmed.ncbi.nlm.nih.gov/22312433/ | Yes, from P14 | Regulon 4$^{th}$ state TF in RGL and GCmat | Candidate NDD gene |
| Klf9 | Delayed maturation of GCs during development + impaired differentiation/decreased neurogenesis in adult DG https://pubmed.ncbi.nlm.nih.gov/19657039/ | Yes, adult brain | No | No results |

| | | | | |
|---|---|---|---|---|
| Atf6 | No | Yes, from P14 | TF in GCmat | No results |
| Prdm5 | No | Yes, adult brain | No | No results |
| Tef | No | Yes, from P14 | Regulon 4$^{th}$ state<br>TF in RGL and GCmat | No results |
| Nr3c1 | Expression decreased in DG upon acute stress https://pubmed.ncbi.nlm.nih.gov/27054829/<br>Lower methylation of Nr3c1 promoter in DG of patients of major depressive disorder https://pubmed.ncbi.nlm.nih.gov/24465557/ | Yes, adult brain | Regulon 4$^{th}$ state<br>TF in RGL and GCmat | No results |
| Cebpd | Alterations in memory and morphology of DG neurons in Cepbd null mice exposed to ionizing radiation https://pubmed.ncbi.nlm.nih.gov/30781689/ | Yes, from P4 | Regulon 4$^{th}$ state<br>TF in GCmat | No results |
| Hlf | Overexpression of Hlf in vivo leads to altered expression of epilepsy-related genes + downregulated in epilepsy models https://pubmed.ncbi.nlm.nih.gov/32111960/ | Yes, from P14 | Regulon 4$^{th}$ state<br>TF in RGL and GCmat | No results |
| Smad3 | LTP impairment in DG upon Smad3 loss https://pubmed.ncbi.nlm.nih.gov/26826552/ | Yes, adult brain | TF in GCmat | No results |

**Table S1. Transcription factors (TF) from SCENIC+ analysis:** expression in the dentate gyrus (DG), link to adult neurogenesis and neurodevelopmental disorders (NDD).

Table S2

| Chemical | Final concentration | Identifier |
|---|---|---|
| **Artificial cerebrospinal fluid (ACSF 1X)** | | |
| NaCl | 126 mM | Sigma S9888 |
| KCl | 2.5 mM | Sigma P3911 |
| NaH2PO4 * H2O | 1.2 mM | Sigma S9638 |
| MgCl2 * 6 H2O | 1.2 mM | Sigma M2670 |
| CaCl2 | 2.1 mM | Sigma C5670 |
| Glucose | 2.7 mM | Sigma G8270 |
| NaHCO3 | 6.25 mM | Sigma S6014 |
| **Digestion Medium (10X DM)** | | |
| MgCl2*6H2O | 27 mM | Sigma M2670 |
| HEPES (1M) | 3.2 ml | Life Technologies 15630056 |
| NaOH (0.1M) | Add until pH=7.35 | Sigma S8045 |
| Na2SO4 | 900 mM | Sigma S6547 |
| K2SO4 | 300 mM | Sigma P9458 |
| ddH2O | top to 200ml | |
| **Digestion Medium (1X DM)** | | |
| 10X DM | 1:10 1X DM | |
| Glucose | 360 µl of 45% Glucose | Sigma G8270 |
| Mg/Kyn stock | 1:10 of Mg/Kyn stock | |
| AP5 (1 mg/ml) | 1:100 | Sigma A5282 |
| PenStrep (10000 U/ml) | 1:100 | Life Technologies 15140122 |
| **Mg/Kyn stock** | | |
| Kynurenic acid | 8 µM | Sigma K3375 |
| MgCl2*6H2O | 37 mM | Sigma M2670 |
| HEPES (1M) | 800 µl (to pH 7.35) | Life Technologies 15630056 |

| | | |
|---|---|---|
| ddH2O | top to 200ml | |
| **OptiMEM/trehalose** | | |
| Optimem | 22.5 ml | Gibco 11058021 |
| Trehalose | 132 mM | Sigma T0167 |
| 45% Glucose | 380 µl of 45% Glucose | Sigma G8270 |
| AP5 (1mg/ml) | 1:200 | Sigma A5282 |
| Mg/Kyn stock | 1:20 | |
| **20% Percoll** | | |
| Percoll working solution | 2ml | |
| Trehalose | 50 mM | Sigma T0167 |
| Optimem | top to 10ml | Gibco 11058021 |
| **Percoll working solution** | | |
| Percoll | 45 ml | Sigma P4937 |
| NaCl 1.5M | 5 ml | Sigma S9888 |

**Table S2. Overview of buffers used for single cell RNA sequencing experiments.** For each chemical compound, final concentration or volume are listed, and commercial brand and identifier.

| RNAscope probes | | | |
|---|---|---|---|
| **Source** | **Identifier** | **Probe** | |
| Bio-Techne | 455311-C1 | RNAscope® Probe - Mm-Bcl6 - Mus musculus B cell leukemia/lymphoma 6 (Bcl6) mRNA | |
| Bio-Techne | 457881-C2 | RNAscope Probe - Mm- Smad3-C2 | |
| Bio-Techne | 317041-C3 | RNAscope® Probe - tdTomato-C3 - tdTomato 1431 bp | |
| **Antibodies IHC** | | | |
| **Source** | **Identifier** | **Antibody** | **Dilution** |
| Aves | GFP-1010 | Chicken Anti-GFP | 1:1000 |
| Takara | 632496 | Rabbit Anti-dsRed | 1:1000 |
| Cell Signaling | 9523 | Rabbit anti-SMAD3 (C67H9) | 1:1000 |
| Roche | ROAHAHA | Rat anti-HA (3F10) | 1:1000 |
| Sigma | A2228 | Mouse anti- β-Actin | 1:5000 |
| Thermofisher | A-11039 | Goat anti-chicken 488 | 1:1000 |
| Thermofisher | A-21428 | Goat anti-rabbit 555 | 1:1000 |
| Thermofisher | 65-6120 | Goat anti-rabbit HRP | 1:5000 |
| Thermofisher | A10549 | Goat anti-rat HRP | 1:5000 |
| Thermofisher | 62-6520 | Goat anti-mouse HRP | 1:5000 |

**Table S3. RNAscope probes and antibodies.** For RNAscope probes, source and identifier are listed. For antibodies, source, identifier and dilution are listed.

### Table S4

| **qPCR primers** | | |
|---|---|---|
| **Primer** | **Sequence** | **Origin** |
| Gfap fwd | CGCTGGCAGCTGAACTGAACCA | In house designed with Benchling |
| Gfap rev | GCACTGTTGGCCGTAAGCTGGT | |
| Prox1 fwd | TGAGCACCTGAGAGCAAAGCGC | |
| Prox1 rev | TCTCGGGGACTCACAGACTGCG | |
| Gphn fwd | TCGCCCAGAATACCACCGGTGT | |
| Gphn rev | TTGGCACTGCGCATGCTCATCA | |
| Slc32a1 (Vgat) fwd | TGAACTGGACACACATCGCCGC | |
| Slc32a1 (Vgat) rev | GCCGGGCAGGTTATCCGTGATG | |
| Calb1 fwd | CAAACTAGCCGCTGCACCACGA | |
| Calb1 rev | CCGTCAGCGTCGAAATGAAGCCA | |
| Aif1 (Iba1) fwd | CTGATGTGGTCTGCACAGGGCG | |
| Aif1 (Iba1) rev | ACAGGCAGCTGAGGAGGACTGG | |
| Pgk1 (HKG) fwd | ACTGTGGCCTCTGGTATACCTG | Calafate et al, Nat Neurosci 2023 |
| Pgk1 (HKG) rev | CAATCTGCTTAGCTCGACCCAC | |
| Bcl6 fwd | AGGGGTTTTGCATCCTCCTGGA | In house designed with Benchling |
| Bcl6 rev | GCTCCATCTGCAGGTACATGGC | |
| Rplp0 (HKG) fwd | AGATTCGGGATATGCTGTTGGC | Eissa et al, PLOS 2016 |
| Rplp0 (HKG) rev | TCGGGTCCTAGACCAGTGTTC | |
| **Genotyping PCR primers** | | |
| **Primer** | **Sequence** | **Origin** |
| Rpb4-Cre fwd | GAACCTGATGGACATGTTCAGG | In house designed with Benchling |
| Rpb4-Cre rev | AGTGCGTTCGAACGCTAGAGCCTGT | |
| RiboTag fwd | GGGAGGCTTGCTGGATATG | JAX Protocol 28336 |
| RiboTag rev | TTTCCAGACACAGGCTAAGTACAC | |
| Ai9:wt fwd | AAGGGAGCTGCAGTGGAGTA | JAX Protocol 29436 |
| Ai9:wt rev | CCGAAAATCTGTGGGAAGTC | |
| Ai9:mut fwd | GGCATTAAAGCAGCGTATCC | |
| Ai9:mut rev | CTGTTCCTGTACGGCATGG | |

| | | |
|---|---|---|
| Rosa26-LSL-Cas9:mut fwd | TGAGCGACATCCTGAGAGTG | JAX Protocol 21206 |
| Rosa26-LSL-Cas9:mut rev | GAGAGCTTTCAGCAGGGTCA | |
| Rosa26-LSL-Cas9:wt fwd | AAGGGAGCTGCAGTGGAGTA | |
| Rosa26-LSL-Cas9:rev fwd | CCGAAAATCTGTGGGAAGTC | |
| H11-Cas9:wt fwd | AGTGGGACTGCTTTTTCCAG | JAX Protocol 29279 |
| H11-Cas9:wt rev | GATCTGGGGCCATAAATGC | |
| H11-Cas9:mut fwd | GGGCAACGTGCTGGTTATTG | |
| H11-Cas9:mut rev | CCAGGCCGATGCTGTACTTC | |

**Table S4. Primer sequences and origin.** RT-qPCR primers and genotyping forward (fwd) and reverse (rev) primers, their sequences and source are listed.

| shRNA or gRNA | Sequence | Source |
|---|---|---|
| shRNA1 Bcl6 fwd | GACACGGATCTGAGAATCT | Tiberi et al, Nat Neurosci 2012 |
| shRNA1 Bcl6 rev | GACACGGATCTGAGAATCT | |
| shRNA2 Bcl6 fwd | TGATGTTCTTCTCAACCTTAA | |
| shRNA2 Bcl6 rev | TGATGTTCTTCTCAACCTTAA | |
| shRNA Ctrl fwd | ACTACCGTTGTTATAGGTG | |
| shRNA Ctrl rev | ACTACCGTTGTTATAGGTG | |
| gRNA1 Smad3 fwd | CCATGGCCCGTAATTCATGG | In house designed with Benchling |
| gRNA1 Smad3 rev | CCATGAATTACGGGCCATGG | |
| gRNA2 Smad3 fwd | GCTGCAGGTGTCCCATCGGA | |
| gRNA2 Smad3 rev | TCCGATGGGACACCTGCAGC | |
| gRNA lacZ fwd | ACCGATGTCGGTTTCCGCGAGGTG | Platt et al, Cell 2015 |
| gRNA lacZ rev | AACCACCTCGCGGAAACCGACATC | |

**Table S5. Short hairpin (sh) and guide (g)RNAs sequences.** Forward (fwd) and reverse (rev) primers used to create shRNA and gRNA, their sequences and sources are listed.
